# Emergence of stable coexistence in a complex microbial community through metabolic cooperation and spatio-temporal niche partitioning

**DOI:** 10.1101/541870

**Authors:** Sonja Blasche, Yongkyu Kim, Ruben Mars, Eleni Kafkia, Maria Maansson, Daniel Machado, Bas Teusink, Jens Nielsen, Vladimir Benes, Rute Neves, Uwe Sauer, Kiran Raosaheb Patil

**Author notes:** These authors contributed equally to the work (Sonja Blasche, Yongkyu Kim).

## Abstract

Microbial communities in nature often feature complex compositional dynamics yet also stable coexistence of diverse species. The mechanistic underpinnings of such dynamic stability remain unclear as system-wide studies have been limited to small engineered communities or synthetic assemblies. Here we show how kefir, a natural milk-fermenting community, realizes stable coexistence through spatio-temporal orchestration of species and metabolite dynamics. During milk fermentation, kefir grains (a polysaccharide matrix synthesized by kefir microbes) grow in mass but remain unchanged in composition. In contrast, the milk is colonized in a dynamic fashion with early members opening metabolic niches for the followers. Through large-scale mapping of metabolic preferences and inter-species interactions, we show how microbes poorly suited for milk survive in, and even dominate the community through metabolic cooperation and uneven partitioning between the grain and the liquid phase. Overall, our findings reveal how spatio-temporal dynamics promote stable coexistence and have implications for deciphering and modulating complex microbial ecosystems.

## Introduction

Complex microbial communities like gut microbiota are characterized by resilience and long-term coexistence (Faith et al., 2013; Mueller et al., 2016; Palleja et al., 2018). On the other hand, ecological models have predicted diminishing stability with increasing community size (Allesina and Tang, 2012; Fyodorov and Khoruzhenko, 2016). Laboratory studies therefore remain crucial for understanding the mechanistic basis of stable coexistence. Experiments with synthetic assemblies or communities with genetically engineered members have helped in, for example, providing empirical evidence for stability at higher species richness (Pennekamp et al., 2018), discover assembly rules (Friedman et al., 2017), and, more mechanistically, in discovering stabilizing interactions like cross-feeding (Blasche et al., 2017a; Dubey and Ben-Yehuda, 2011; Ponomarova et al., 2017; Wintermute and Silver, 2010). These interactions allow, for example, collective exploitation of resources more efficiently than any single species on its own (Alessi et al., 2018; Rosenthal et al., 2011). Furthermore, environmental changes can favor certain species or maintain diversity by temporally partitioning the growth of different species (Baran et al., 2015; Ratzke and Gore, 2018; Yuan and Chesson, 2015). In large communities, both inter-species and species-environment interactions often act simultaneously leading to fascinating but intricate dynamics. This complexity has been difficult to dissect experimentally as the current experimental communities consist of few species, are cultivated *ex situ* under different conditions, or require inoculation from synthetic assemblies or original sources (Embree et al., 2014; Ponomarova et al., 2017; Venturelli et al., 2018; Wolfe et al., 2014). How natural communities realize stable coexistence has thus far remained mainly a question for theory.

Here, we experimentally tackle this question using kefir as a model microbial ecosystem. Kefir has suggested origins in the Caucasian mountain region (Bourrie et al., 2016; Motaghi et al., 1997); with an early evidence suggesting kefir usage from an artefact on a circa 3,500 years old Chinese mummy (Yang et al., 2014). The kefir community features a rich diversity encompassing approximately 30-50 species (Prado et al., 2015), including both prokaryotes (predominantly lactic and acetic acid bacteria) and eukaryotes (yeasts) (Figure 1A). Despite this rather large membership, cultivation and maintenance of the community is straightforward as the kefir grains consisting of extracellular polymeric substances, act as a self-renewing community-scale inoculum. Furthermore, the kefir community is highly resilient to various abiotic as well as biotic stresses such as desiccation and non-sterile handling (Farnworth, 2008). These features collectively motivated our choice of kefir as a model ecosystem.

**Figure 1.**
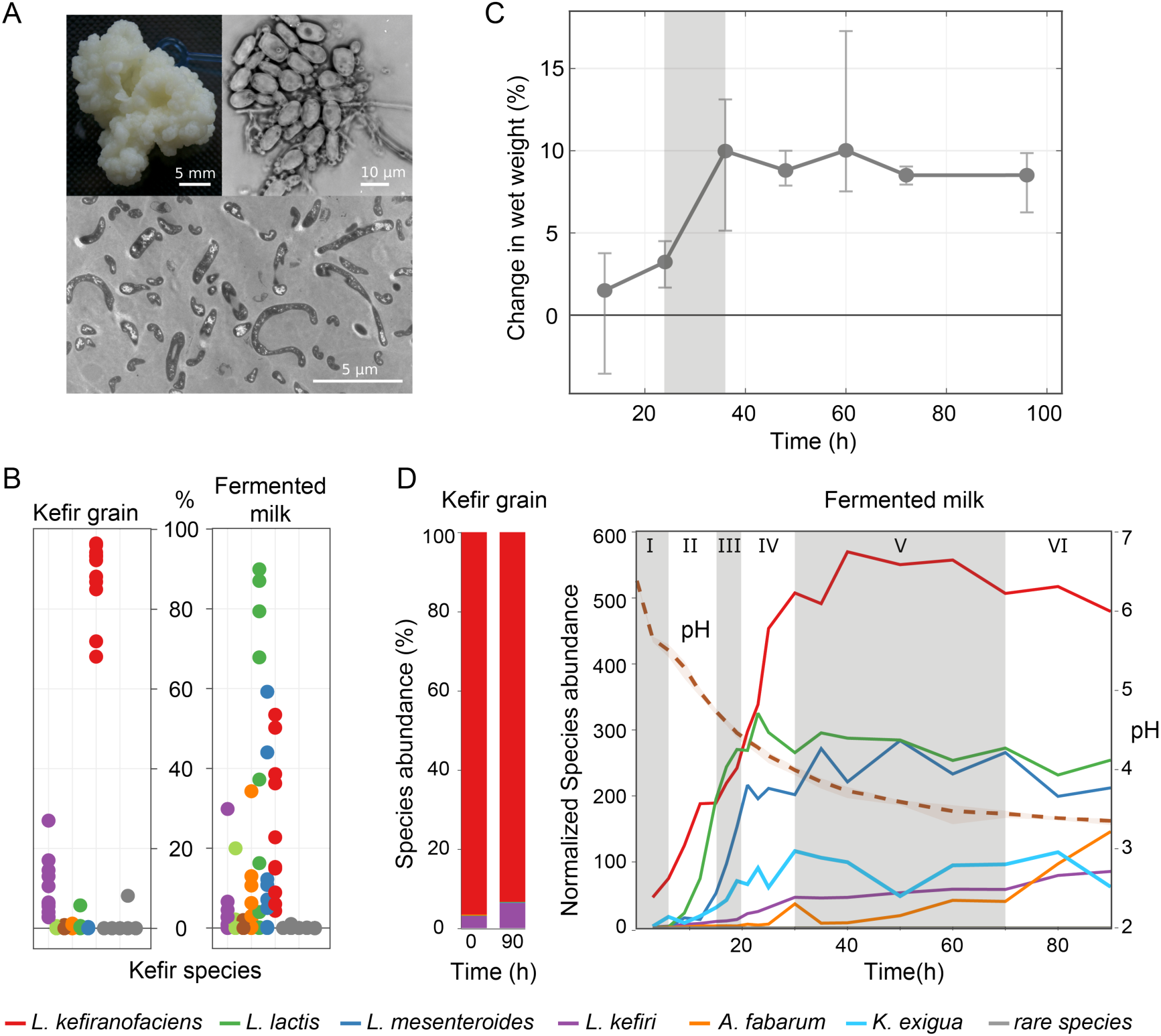
Diversity and dynamics of kefir communities. (A) Example images of kefir grain and its associated microbial community: macroscopic grain (top left), light microscopy image of a region with high yeast density (top right), and a transmission electron microscopy image showing bacteria embedded in the grain matrix. (B) The species composition of different kefir grains and fermented milk fractions. All kefir cultures harbored the same major species and only some of the low abundance species were found to be community specific. (C) Weight gain of the reference kefir grain (kefir GER6, OG2) during milk fermentation. (D) Species dynamics in the grain and fermented milk during kefir fermentation assessed by 16S amplicon sequencing (bacteria) and qPCR (yeast).

## Results

### Kefir grains of different origins harbor a common core set of microorganisms

To choose a particular kefir community for in-depth analysis, we started by profiling the species diversity of kefir grains collected from different geographical locations (6 German, 3 UK, 1 Korean and 1 Turkish, see Table S1: List of kefir cultures) and their fermented milk products, harvested after 48 hours fermentation. Supporting their suspected common origin and in line with the results from previous studies (Gao et al., 2012; Garofalo et al., 2015; Marsh et al., 2013), we found that the vast majority of grains featured a bacterial core community structure consisting of *Lactobacillus kefiranofaciens*, *Lactobacillus kefiri*, *Leuconostoc mesenteroides, Lactococcus lactis*, and *Acetobacter sp.*, which together accounted for more than 95% of total abundance. The grains differed only in the harbored yeasts and low abundance bacteria (rare species) (Figure S1). We also performed shotgun metagenomic sequencing of two selected grains (different origins), which further confirmed these findings (see Table S2: Metagenome sequencing table, kefir GER6). Interestingly, the bacterial composition between the solid (grain) and the liquid (milk) fractions were strikingly different in all cases (Figure 1B and Figure S2), suggesting that the community undergoes complex dynamics during fermentation.

### Kefir fermentation does not change the grain composition but involves sequential colonization of milk

To gain insights into the dynamics of the kefir community, we profiled, using one of the kefir grains collected in Germany (referred to as GER6), grain and species growth over a 90 hours long fermentation. For the milk fraction, 18 time points were chosen based on pH dynamics, sampling more densely during the early hours when pH dropped rapidly. The grains were analyzed in the beginning and at the end. Bacterial and yeast composition was monitored using 16S rDNA and ITS amplicons, respectively. The resulting relative abundance data was then scaled to absolute values using qPCR analysis for yeast (*Kazachstania exigua*) and *L. kefiranofaciens* in combination with the total DNA.

The kefir grain followed a sigmoidal growth pattern with a short lag phase in the beginning, followed by a steep increase in weight until it reached a plateau at 36 hours (Figure 1C). The 16S profiles showed that the species composition in the grain remains constant, whereas that in the fermented milk fraction underwent remarkable changes, particularly in the first 30 hours of fermentation (Figure 1D, Figure S3). Based on these compositional dynamics of the milk fraction, we divided the fermentation into six phases. Phase I is a lag phase where *L. kefiranofaciens*, dominant in the grain, is most abundant as expected. Phase II is marked by rapid growth of *L. lactis*, followed by phase III wherein *L. mesenteroides* is the fastest growing species. *L. kefiranofaciens*, which showed continued growth throughout the initial three phases, reached its stationary phase near the end of the phase IV. This phase is also the initial time frame for the growth of *L. kefiri*, which increased to be the fifth most abundant member species in the community. Growth of all species, except *Acetobacter* stagnates in phase V, which is followed by the final phase (VI) wherein *A. fabarum* and *L. kefiri* are the only appreciably growing species. The abundance of the eukaryotic member of the community, *K. exigua* increases gradually up to phase IV, reaching a stable plateau after 30 hours of fermentation.

### Metabolomics reveals complex nutrient and by-product dynamics during kefir fermentation

To elucidate the metabolic changes accompanying the species dynamics observed during kefir fermentation, extracellular metabolites were monitored using different untargeted and targeted methods. Untargeted analysis was done by using flow injection analysis time-of-flight mass spectrometry (FIA-qTOF) (Fuhrer et al., 2011) (see Methods). This revealed extensive metabolic changes as fermentation proceeds, some of which suggest sequential niche creation (Figure 2A). Further tracking by unsupervised K-means clustering, revealed six clusters (Figure S4) that captured different patterns of metabolite changes (Figure 2B). In particular, the inflection points, i.e. the time points at which the rate of metabolite change shifted in magnitude or direction, coincided with the distinct growth phases derived from the species dynamics (Figure 2B and Figure S5). This correlation suggests a major role of species activity on metabolite dynamics, which may, in turn, determine time windows of opportunity for different community members. The first two clusters group metabolites accumulating throughout the fermentation and include many amino acids (Table S3: FIA-qTOF k-means_clusters), whereas cluster three represents metabolites that are produced early on and consumed as the fermentation proceeds: mainly sugars and carboxylic acids.

**Figure 2.**
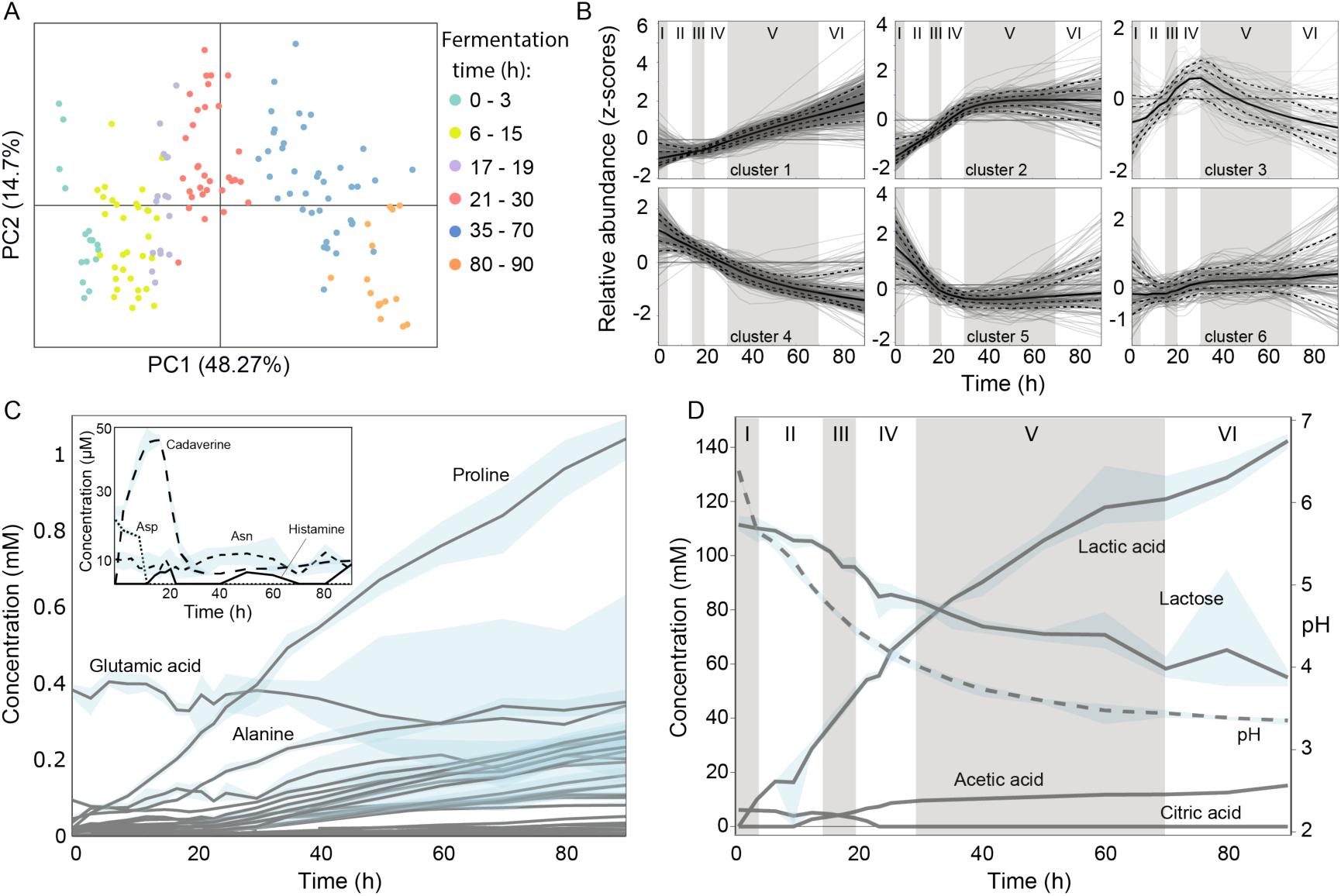
Metabolite dynamics during kefir milk fermentation. (A) PCA analysis of untargeted metabolomics data. Samples are colored according to the six fermentation phases as determined by species dynamics. (B) Temporal dynamics of metabolite ions during fermentation in kefir fermented milk. All the ions detected by FIA-qTOF MS were grouped into 6 clusters by k-means clustering. The solid line represents the median values of the ions in each cluster and the dashed-lines include 10%, 25%, 75% and 90% of the metabolites, respectively. The six fermentation phases are marked by roman numerals. Quantitative assessment of (C) free amino acids and polyamines, and (D) carbohydrates and pH changes during kefir fermentation.

To validate the metabolic changes revealed by the untargeted method and to quantitatively assess the dynamics of major metabolite players, we measured amino acids, polyamines, sugars, and organic acids by using ion chromatography and GC-MS (see Methods). As expected, lactose was consumed throughout the fermentation with corresponding increase in lactate concentration. In the first 20 hours, where the pH drop was most rapid, the overall lactose metabolism shifted from homo-fermentative to hetero-fermentative mode as reflected in the lactate yield on lactose (Figure S6). This was in good concordance with the sequential growth of homo-fermentative (*L. lactis* and *L. kefiranofaciens*) and hetero-fermentative lactic acid bacteria (*L. mesenteroides*). Yet, about half of the initial lactose was left after 90 hours when the pH plateaued at 3.35 (Figure 2D). Also, no appreciable histamine production was observed, in contrast to other fermented milk products (Linares et al., 2012). Consistent with the untargeted analysis (clusters 1 and 2 in Figure 2B), most amino acids accumulated throughout the fermentation (Figure 2C), implying that the sustained proteolysis of milk protein by the bacteria surpasses their growth demand. Exceptions were asparagine and glutamate whose concentrations remained at initial levels throughout and aspartate, which was rapidly consumed in the first few hours and thereafter remained undetectable. For aspartate and glutamate this pattern is in stark contrast with the milk total protein composition and both were predicted by genome-scale metabolic modeling to be substantially consumed by kefir species (Supplementary figure S7). Aspartate and glutamate thus stand out as the main requirements of kefir species.

### Citrate depletion kickstarts the community growth

The total amino acid accumulation markedly increased after around 20 hours, which coincides with the depletion of milk citrate (Figure 2D, Figure S8). We hypothesized that this is due to the metal ion chelating property of citrate (Apelblat, 2014) that inhibits metal dependent proteases secreted by lactic acid bacteria such as *L. lactis* (Law and Haandrikman, 1997). Supporting this, we observed that the growth of the grain is dependent on proteolysis and can be inhibited by adding another metal ion chelator, EDTA (Figure S9), which also inhibits growth of several kefir species (Figure S10). Further, grain growth in kefir spent whey harvested after 36 hours, in which its growth is strongly impaired, can be restored by the addition of casein hydrolysate. On top addition of EDTA eliminates this effect (Figure S9) indicating a connection between proteolysis and community growth. Removal of citrate from the milk is well correlated with the early-growers *L. lactis* and *L. mesenteroides* that are known citrate utilizers (Marty-Teysset et al., 1996; Samaržija et al., 2001). Citrate removal thus appears to be one of the first metabolic functions crucial for kickstarting proteolysis and thereby kefir grain growth.

### Inter-species dependencies are essential for the survival of kefir species

Towards disentangling the role of different species and their interaction with the dynamic environment in the fermenting milk, we isolated 40 different strains of bacteria and yeast from kefir. These spanned all abundant species as well as several rare species like *Rothia dentocariosa*, together representing ~99 % of the species hitherto found in their source kefir community (Table S4: List of isolated kefir species). All isolates were genome sequenced and four were found to represent completely novel species (Blasche et al., 2017b; Kim et al., 2017). To assess the milk colonization capability of the individual strains, we measured their growth in milk whey (Figure S11, Table S5 “Species OD kefir+milk whey”). Strikingly, only yeasts and some low abundance species, but none of the more abundant kefir bacteria, showed high growth in this medium in monoculture. A notable example is *L. kefiranofaciens*, which is monodominant in the grain and top abundant in the milk fraction, but did not show considerable growth either in milk whey nor in milk (Figure S12). *L. kefiranofaciens* thus appears to be completely reliant on its fellow community members for survival.

The growth support for a species, e.g. in terms of essential nutrients, could either come from the activity of the previous members or may in addition require concomitant presence of another species. We first assessed the extent to which the metabolic activity of the prior members provided the necessary growth conditions (niche opening). For this, we grew the isolated kefir species and also few selected non-kefir species in kefir spent whey prepared at different fermentation stages from 12 to 96 hours (Figure 3A, Figure S11 and Table S5 “Species OD kefir+milk whey”). The kefir spent whey is in essence a medium conditioned by the kefir community including all soluble metabolites present at the sampling time. The growth preferences of four of the five most abundant bacterial species in the kefir spent whey matched well with their growth time window during the fermentation: *L. kefiranofaciens* and *L. mesenteroides* are early-growers, *Acetobacter fabarum* only commences growth after 36 hours and *L. kefiri* steadily increases during the whole fermentation course. The latter two species thus seem to obtain their resources mainly through the metabolic activity of the preceding bacteria. In contrast, none of the *L. lactis* strains, including non-kefir variants, showed the early stage growth preference observed in kefir fermentation, and even showed impaired growth in the kefir spent whey, suggesting need for concomitant presence of other species or missing nutrients in the whey. Interestingly, *L. kefiranofaciens* did not grow alone neither in milk nor in milk whey, but grew in early harvested kefir whey (Figure S12), suggesting that it is supported by another early grower species. Together, the growth patterns in the milk and the kefir whey brought forward the crucial role of inter-species dependencies in the survival of the key kefir species and that of the overall community.

**Figure 3.**
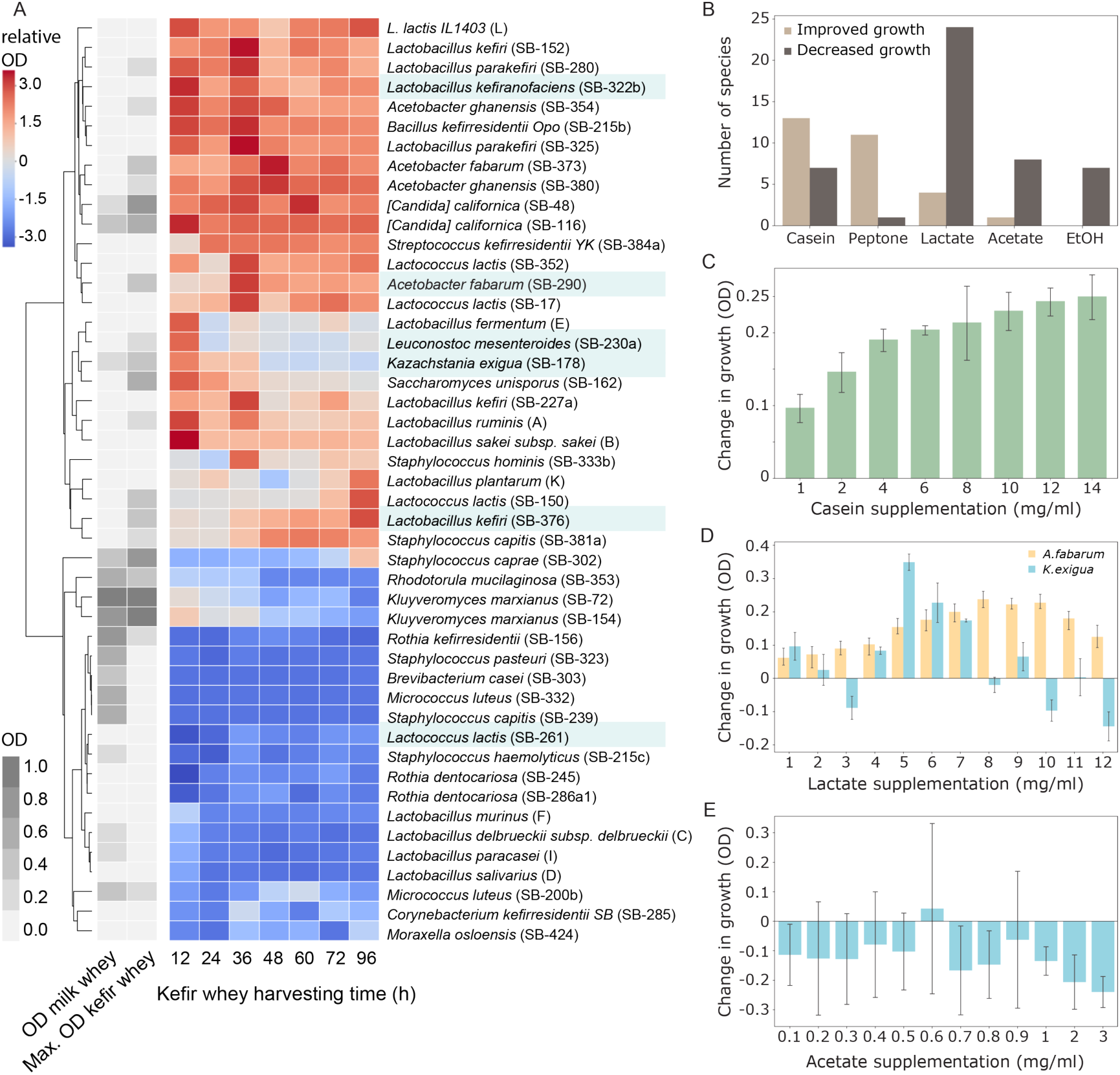
Growth performance of kefir isolates. (A) Growth of kefir isolates in milk whey and kefir whey (i.e. community spent medium) collected at different fermentation stages. For kefir whey, growth measurements (OD) relative to that in the milk whey are shown (relative OD). Main kefir species are highlighted. (B) Growth impact of major fermentation products, viz., lactate, acetate and ethanol, and of supplemented protein sources casein and peptone added in different concentrations to milk whey. Species displaying significant changes (t-test, p<0.01) in at least 70% of the different concentrations were counted as being promoted or suppressed by the tested component. (C) Effect of casein on growth of *L. lactis*. During whey preparation, most of the casein is removed, so that milk whey only contains about 20% of the protein content of bovine milk. Protein supplementation of milk whey with 1 to 16 mg/ml casein hydrolysate partially restored natural protein content. (D) Lactate supports *Acetobacter* and *K. exigua* growth in milk whey at different concentrations. (E) Effect of acetate on growth of *K. exigua.* Changes in species growth are assessed relative to growth in non-supplemented milk whey.

### Major by-products of milk fermentation determine the growth window for yeast and *Acetobacter*

The stark difference between the milk and kefir spent whey growth suggested that the major fermentation products play an important role in determining the growth windows of different species. To test this, we re-evaluated the growth of individual strains in the milk whey, but now supplemented with lactate, acetate or ethanol. In addition, we also tested the effect of casein hydrolysate (and peptone) supplementation to account for the protein removed during whey production. Overall several rare species, and especially *L. lactis*, profited from protein supplementation and were inhibited by acetate and lactate (Figure 3B-C, Figure S13). Lactate was particularly inhibiting for low abundance species. Lactate and acetate also benefited certain species, albeit in a concentration-dependent manner. *Acetobacter sp.* benefitted from lactate at concentrations between 7 and 10 mg/ml that are reached after 35 hours of kefir fermentation. *Kazachstania exigua* (and *Saccharomyces unisporus*) also benefited from lactate but at lower concentrations (Figure 3D, Figure S13). Acetate, on the other hand, inhibited kefir yeasts, esp. *K. exigua* (Figure 3E), and the effect differed from that observed for Sa*ccharomyces sp.* and *Kluyveromyces marxianus* that gained profit from acetate concentrations below the inhibitory amount of 2 mg/ml (Figure S14) (Greetham, 2015; Wright et al., 2011). Indeed, the window of yeast growth and the start of the *Acetobacter* growth during kefir fermentation could be explained with the balance between the positive and negative effect of lactate and acetate on these species (Figure S15).

### Species interaction network

We next set out to identify particular interactions between different kefir species. To capture different modes of interactions, we used three distinct readouts assessing the performance of binary communities against the corresponding single strains: i) milk acidification rate, indicating overall fermentation competence (Figure 4A); ii) growth in milk (quantified by 16S amplicon sequencing relative to a spiked-in *E. coli* standard) (Figure 4C); and iii) colony size on milk-based agar plates (Figure 4F).

**Figure 4.**
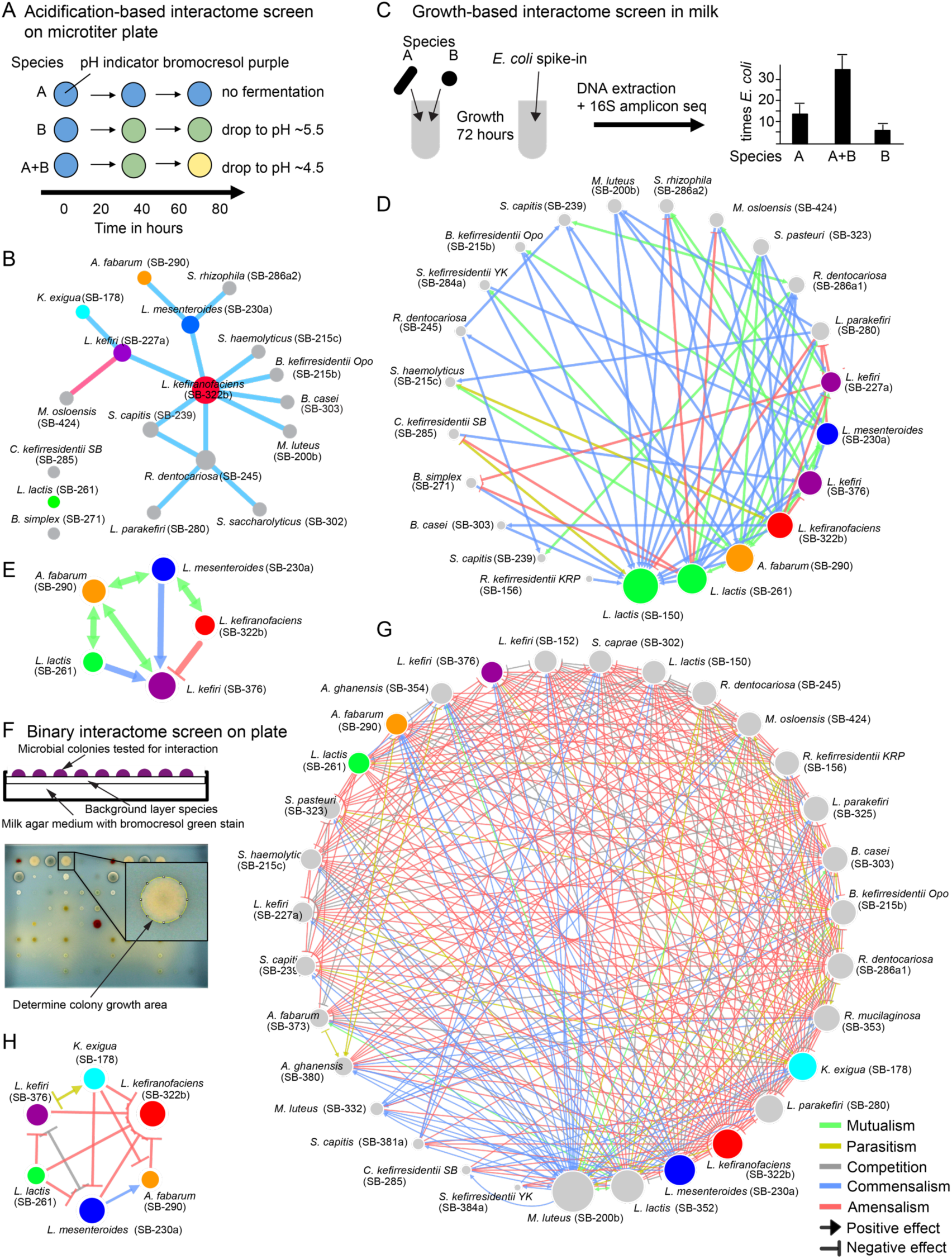
Kefir species interaction networks mapped using different approaches. (A) Schematic depiction of the method used to map metabolic interactions based on fermentation kinetics. Species were grown in 96-well plates alone or in pairs; acidification of milk was assessed with a soluble pH-indicator. Positive interactions were those that increased acidification in coculture compared to single species, while negative interactions showed decreased acidification. (B) Network of metabolic interactions between kefir species mapped using (A). (C) Schematic description of the method used to assess growth promoting or inhibiting interactions between kefir species in milk. Species abundance in mono- and cocultures was assessed by 16S amplicon sequencing with quantitative *E. coli* spike-in. The interactions thus revealed are shown in (D) for all tested kefir species, and in (E) for the major species in milk. (F) Method used for detecting species interactions on milk plates. Each of the species was plated as a background layer on milk plates containing bromocresol green. All query species were pinned on top and the colony area was assessed as a growth metric. (G) and (H) depict the resulting interaction networks between all tested and major kefir species, respectively. Node sizes represent the number of interactions found for the respective species.

The acidification-based interaction assay revealed a significant interaction (p<0.05) for 15 cocultured pairs (Figure 4B, Table S6: Acidification-based interactome). *L. lactis* did not show any interactions and was able to ferment on its own as well as in company, consistent with its early growth behavior during the kefir fermentation. In contrast, eight of the interacting pairs included *L. kefiranofaciens*, the dominant species in the grain as well as in the fermented milk fraction. While only two of those interactions involved more abundant lactic acid bacteria, *L. mesenteroides* and *L. kefiri*, six were with rare species. Furthermore, we observed that some of these rare species supported the growth of the kefir grain when added to the milk before starting the fermentation (Figure S16). Combined with the high number of interactions between rare species and *L. kefiranofaciens*, this points to their functional role and evolutionary selection for inclusion in the kefir community.

Though functionally important, the acidification assay could not reveal whether only the fermentation activity was altered during the coculturing, or whether the growth of the individual microbes was also changed. We therefore monitored growth changes by using a 16S amplicon sequencing-based approach with quantification relative to a spiked-in *E. coli* standard. This revealed 70 interactions: 58 positive (mutualism or commensalism) and 12 negative (competition, amensalism, or parasitism) (Figure 4D, Table S7: Growth-based interactome-milk). When only considering interactions between the abundant kefir species (Figure 4E), *L. kefiranofaciens* suppressed its seemingly direct competitor *L. kefiri*, while promoting growth of *L. mesenteroides* and having no effect on *L. lactis* and *A. fabarum*. The majority of the mutualistic interactions, 8 in total, involved *A. fabarum* and a lactate-producing partner, supporting its dependency on lactate during the late onset of its growth during the kefir fermentation.

To assess interactions in a solid medium, which is better mimicking the conditions in the grain, we tested all pairs of kefir isolates and selected non-kefir *Lactobacilli* (Table S8: Non-kefir lactic acid bacteria) (in total 48 strains) for interaction on milk-agar plates. A given strain (background) was plated as a lawn and all others (queries) were pinned on top. Change in the colony size relative to “no background” was used to call interactions (Figure 4F, Methods). This revealed a dense network with strikingly higher number of negative interactions than observed in the liquid medium, both among major kefir species and in the whole network (Figure 4G-H, Table S9: Growth-based interactome-plate). While 42.8 % interactions in the liquid milk medium were positive and only 7.4 % negative, the solid medium featured only 8.3 % positive but 53.9 % negative interactions (Figure S17). Especially *L. lactis*, known to produce bacteriocins was a potent inhibitor of other species (Figure S18). Thus, while kefir species seem to support each other in the liquid phase, competition was predominant on the solid medium.

The stark reversal of the cooperation-competition divide between the solid and the liquid medium provide an explanation for the similarly stark difference in the community composition between these two phases. The grain is monodominated by *L. kefiranofaciens*, which is the main producer of the grain matrix and thus has a preferential access to this niche. It experiences mostly inhibitory interactions from the lower abundant species that thrive to maintain themselves in the grain. In contrast, thriving in the liquid milk requires cooperation and thus this niche is more dynamic and the diversity is more evenly distributed.

### Amino acids and lactate drive the mutualistic interaction between *L. mesenteroides* and *L. kefiranofaciens*

The mutualistic interaction between *L. kefiranofaciens* and *L. mesenteroides*, observed in the milk medium, is an early event during kefir fermentation and thus a likely determinant of subsequent species dynamics. To elucidate the factors mediating this interaction (Figure 5A), we searched for metabolites/factors that could compensate for the interacting partner in terms of growth promotion. We therefore supplemented milk with different combinations of amino acids, mineral mix, vitamin mix, DNA bases (including uracil), DNAse treated salmon sperm DNA, xanthine, NAD, and inactivated inoculum (either *L. kefiranofaciens* or *L. mesenteroides* killed by ethanol or heat inactivation, supplied in 1x or 10x amounts) (see Methods). We then monitored milk acidification of the *L. kefiranofaciens* or *L. mesenteroides* monocultures over 76 hours. None of the supplements boosted fermentation of *L. kefiranofaciens* as observed in coculture with *L. mesenteroides*. However, a combination of trace minerals, vitamins and amino acids restored the fermentation profile of *L. mesenteroides* to a similar degree as coculture with *L. kefiranofaciens* (Figure 5B).

**Figure 5.**
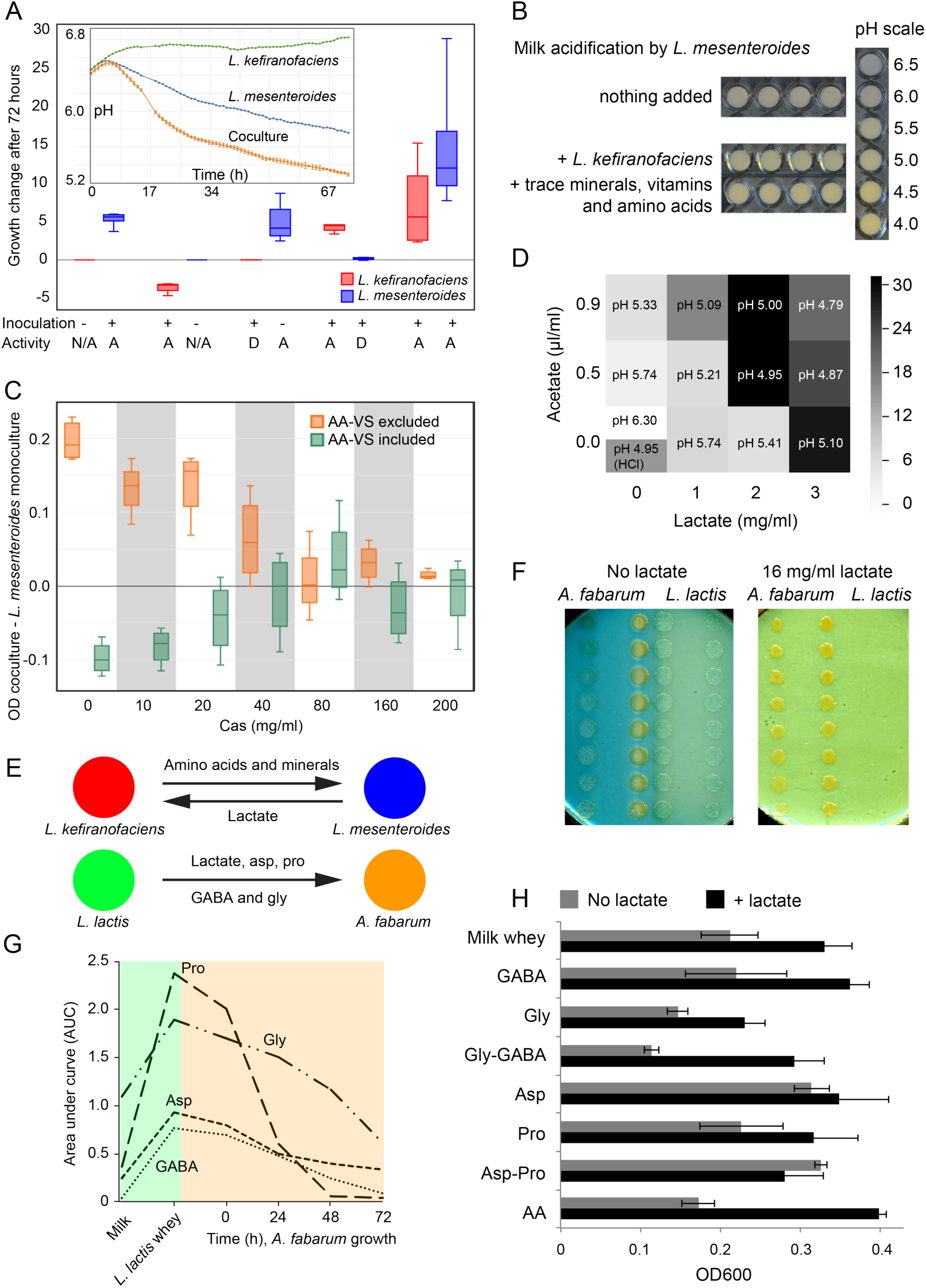
Unraveling species interactions. (A) Interaction between *L. kefiranofaciens* and *L. mesenteroides. L. kefiranofaciens* and *L. mesenteroides* profit from each other when both are alive. Whereas *L. kefiranofaciens* receives similar benefit from dead *L. mesenteroides*, *L. mesenteroides* only profits from *L. kefiranofaciens* when alive; D=dead (heat or ethanol treated), A=alive, N/A=not present. (B) Addition of amino acid mix, vitamin mix and trace minerals to milk improve acidification by *L. mesenteroides* after 72 hours fermentation in a way comparable to the on top addition of *L. kefiranofaciens*. (C) *L. mesenteroides* growth (OD600) in monoculture compared to its coculture with *L. kefiranofaciens* in milk whey enriched with amino acids (AA), vitamins (VS) and different amounts of casein peptides (Cas, in mg/ml). Up to a supplementation of 20 mg/ml casein, mono-and cocultures grow significantly differently (p<0.05). The addition of extra AA and VS boosts monoculture growth above that of the coculture. The effect disappears after supplementation of 40 mg/ml or more casein. (D) Supplementation of lactate and acetate in different proportions support growth of *L. kefiranofaciens* in milk. The effect is a combination between lowered pH (which alone improves growth at pH 4.95, adjusted with HCl) and lactate availability. Acetate supplementation did not exceed the effect of pH alone. (E) Identified interactions between *L. kefiranofaciens* and *L. mesenteroides*, and between *A. fabarum* and *L. lactis*. (F) *A. fabarum* and *L. lactis* in coculture on milk agar plates with and without lactate supplementation (16 mg/ml). (G) Aspartate, proline, glycine and GABA accumulation by *L. lactis* and consumption by *A. fabarum* in *L. lactis* spent whey. (H) *Acetobacter* growth in milk whey, and in milk whey plus 8 mg/ml lactate, supplemented with different amino acids and GABA that it was found to consume. Lactate additionally boosts *Acetobacter* growth in all cases except aspartate (and aspartate plus proline) supplementation, where it grows equally well with and without the lactate supplementation.

To further pinpoint the growth requirements of *L. mesenteroides*, we continued with a milk whey-based growth medium, since this allowed for direct growth measurements. On this medium, we tested the effect of supplementation with free amino acids, vitamins and small peptides (<3 kDa) derived from proteinase K digested casein hydrolysate (further referred to as casein peptides). These experiments showed that either casein peptides or free amino acids could restore the growth of *L. mesenteroides* at the levels observed in the coculture (Figure 5C). Furthermore, active proteinase K boosted its growth in peptide-rich milk whey (Figure S19), underscoring the dependence of *L. mesenteroides* on proteolytic activity by other community members. This is in agreement with a previous study reporting that *L. mesenteroides* requires additional trace metals (Mn^2+^ and Mg^2+^) and amino acids for its growth in milk (Bellengier et al., 1997). Our results, however, show that the trace metals are not required *per se*. However, they could play a role due to metal ion sequestration by milk micelles (Davies and White, 1960). Nevertheless, as we show, proteolytic activity can both provide the needed amino acids as well as overcome the limitation in metal ion availability. Overall, proteolytic activity of *L. kefiranofaciens* constitutes the beneficial effect on *L. mesenteroides* growth.

To uncover the metabolic feedback from *L. mesenteroides* to *L. kefiranofaciens*, we further tested the effect of lactate and acetate supplementation, as these two are major by-products of *L. mesenteroides* metabolism (Hache et al., 1999) and lactate was predicted to be possibly exchanged by model simulation (see Methods). Indeed, lactate could considerably boost (~ three fold) the growth of *L. kefiranofaciens* (Figure 5D). In contrast, acetate supplementation, pH change, and proteolysis, had only a minor contribution to this effect (Figure S20). This concords well with the model prediction that lactate can be uptaken by *L. kefiranofaciens* and converted to acetate yielding one molecule of ATP.

Taken together, the metabolic cooperation between *L. kefiranofaciens* and *L. mesenteroides* is mediated by amino acids and small peptides liberated by proteolytic activity of *L. kefiranofaciens*, which in turn benefits from lactic acid produced by *L. mesenteroides* (Figure 5E).

### *Acetobacter* profits from aspartate and lactic acid

We next investigated the metabolic dependencies and the main benefactor species of *Acetobacter*, which is the latest grower and has low fitness in milk on its own. Since lactate supplementation improved *Acetobacter* growth in milk whey (Supplementary Figure S13), and LAB, esp. *L. lactis*, showed positive effect on its growth (Figure 4), we hypothesized that lactic acid and possibly other factors produced by *L. lactis* were beneficial for *Acetobacter*. Conforming to this, *Acetobacter* grew closer to *L. lactis* colonies on milk plates, an effect that could be recapitulated using lactate addition (Figure 5F). To test whether lactate was consumed, and to discover other potentially exchanged metabolites, we performed metabolomic analysis of time-compartmentalized interaction using a conditioned media set-up. Spent whey prepared from four *L. lactis* strains were used to grow four *Acetobacter* isolates and supernatant samples were collected for subsequent analysis. Untargeted FIA-qTOF analysis showed proline and aspartate exchange between all *L. lactis* and *Acetobacter* strains (Figure S21, Table S10: *L. lactis-Acetobacter* FIA-qTOF). Targeted mass-spectrometry analysis using HILIC-qTRAP and GC-MS confirmed these results and GC-MS further identified two additional exchanged metabolites, glycine and 4-aminobutyric acid (GABA, a neurotransmitter) (Figure 5G, Supplementary figure S22). The latter was recently shown to be produced by *L. lactis* from glutamate (Lacroix et al., 2013). Interestingly, though GC-MS data showed a huge accumulation of lactic acid as expected, no appreciable consumption was observed (Figure S23). This could be due to much smaller need by *Acetobacter* compared to the large availability and thus the consumption would be within the variation of the method. We also observed that glycine, despite being consumed in the conditioned media assay, is also inhibitory (Figure 5H). The consumption of proline from *L. lactis* is unlikely to be limiting in the context of whole community as proline accumulates in large quantities (Figure 2C). Aspartate, on the other hand, is likely limiting during kefir fermentation (Figure 2C) and also boosted the growth of *Acetobacter* in milk whey (Figure 5H). In addition, *Acetobacter* can synthesize aspartate from lactate (Adler et al., 2014), which could also promote its growth. Together, the metabolomics and supplementation results show that *Acetobacter* grows in later stages due to the availability of lactate that it uses as a basis to produce amino acids, especially aspartate, which is released by casein proteolysis but directly consumed by its predecessors in early fermentation stages.

## Discussion

Our data showcases dynamics and inter-species interactions underlying the co-existence in a complex natural community in its native environment and reveal its mechanistic principles. Perhaps the most fascinating aspect is how *L. kefiranofaciens* dominates the community despite having no fitness in milk on its own. This becomes even more striking when considering that kefir is its only known natural habitat. Most other kefir species have been found in other environments. The community thus has likely co-evolved to support *L. kefiranofaciens* while also benefitting from the opportunity to colonize a new niche. The cooperative interactions that we identified, though showing how *L. kefiranofaciens* can survive in milk, do not fully explain its overly large abundance (circa 95% in grain and up to 40% in milk). The key to this lies in the way the community is propagated over time by the use of kefir grains - mainly constructed by *L. kefiranofaciens* - as inoculum. The abundance profile of the community has thus been directly shaped by the grain rather than the milk. Thus, *L. kefiranofaciens* has evolved to maintain its dominance in the grain it builds. The other members of the community, however, must also be carried along as otherwise the growth in the fresh medium would be hampered.

We revealed the role of several community members through extensive analysis of species and metabolite dynamics, characterization of individual members, and charting of inter-species interactions. Though milk is a rich medium, many of its nutrients, such as amino acids, are not readily available and need to be released through proteolysis. An important function of early-growers like *L. kefiranofaciens* and *L. lactis* is to release the amino acids. On the other hand, *L. kefiranofaciens* depends on *L. mesenteroides* for its own growth. By-products of fermentation like lactate are consumed by the late-growers but also act as inhibitors balancing the growth of, e.g., yeast. Indeed, many of the low abundant members of the community, like *Rothia, Staphylococci, Brevibacterium* and *Micrococcus*, are relatively good growers in milk but are quickly outcompeted due to their sensitivity to fermentation products. Nevertheless, these rare species are stably integrated into the community due to their contribution in terms of proteolysis and as they find a time-window of growth, albeit short, during the complex dynamics. These species accordingly showed several positive interactions with their more abundant fellow species. Several negative interactions were also mapped, suggesting that the overall community structure is held through both positive and negative interactions.

Overall, the stable coexistence of the kefir community can be conceptually pictured as a “basecamp model” (Figure 6). The grain is used by the community members as a basecamp, from which the members colonize the milk in an orderly fashion orchestrated by the accompanying metabolite dynamics. The growth of the grain during fermentation recruits the community members for the next transfer. Supporting this model, we observed declining abundance of *L. kefiranofaciens* within a few transfers when the community was passaged without the grain (Figure S24). The spatio-temporal niche separation underlying this basecamp lifestyle might be a key mechanism also in other microbial systems. For example, species retained in the mucosal layer of the intestines can provide a basis for community restoration after dysbiosis events like antibiotic treatment. From an engineering perspective, our results can be used to design new stable communities or to pinpoint the mechanisms underlying destabilization of natural communities and to design appropriate intervention strategies. To this end, kefir provides an excellent model system.

**Figure 6.**
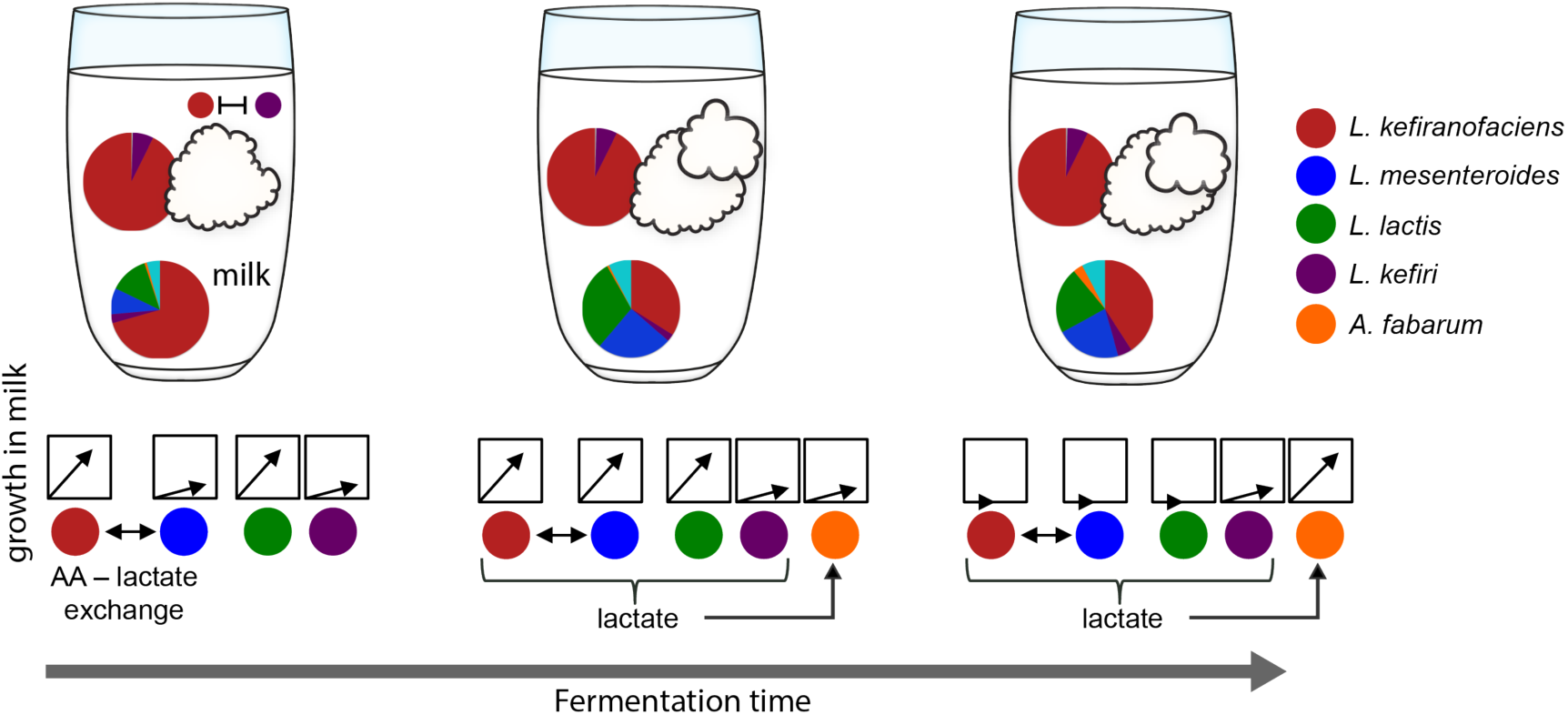
Conceptual model of the kefir community’s “basecamp” lifestyle. The community in the grain itself shows only minor compositional changes, while the milk fraction continuously changes as fermentation proceeds. The grain is thus used by the community members as a basecamp, from which the members colonize milk in an orderly fashion orchestrated by the accompanying metabolite dynamics (see Figure 1,2). The growth of the grain during the fermentation recruits the community members for the next transfer to the fresh milk or storage in-between.

## Author contributions

S.B. and Y.K. conceived the project, performed experiments, analyzed the data and wrote the paper. R.M. performed untargeted metabolomics experiments and data analysis, and quantitative analysis of aspartate and proline. E.K. and M.M. performed GC-MS and ion chromatography analysis. V.B. contributed to the amplicon and metagenome sequencing. J.N and B.T. contributed to experimental design. R.N. oversaw the targeted metabolomics analysis. D.M. contributed to amino acid profile analysis and interpretation. U.S. oversaw the exo-metabolome analysis. K.R.P. conceived the project, designed experimental approach, oversaw the project and wrote the paper.

## Acknowledgements

We thank EMBL Genecore facility for help with (meta-)genomic sequencing, O. Ponomarova for help in collecting kefir grains, advice on cultivation, and for feedback on the manuscript, and K. Zirngibl for feedback on the manuscript. This work was sponsored by the German Ministry of Education and Research (BMBF, Grant number 031A601B) as part of the ERASysAPP project SysMilk.

## Supplemental information

Supplemental Information includes Extended Experimental Procedures, 24 figures, and 13 tables. Genome-scale metabolic models for kefir bacteria can be found at https://github.com/cdanielmachado/kefir_models

## Methods

### Kefir propagation

Before inoculation, kefir grains were washed with double distilled H_2_O and the weight was determined. 60g/l were inoculated in 3.5% UTH cow milk (MUH Arla eG, Germany) and propagated at room temperature without shaking for 48 hours.

**The kefir fermentation curve** was run in quadruplicates for 90 hours (h) total time. Each replicate contained 650ml total volume with 60g/liter kefir grains. PH measurement, DNA and metabolite sample collection was done from 0-15h, 15-25h, 25-40h and 40-90h every 3, 2, 5 and 10 hours, respectively.

**DNA extraction** was done using a two-step approach combining enzymatic digestion and bead beating (modified after (Kowalczyk et al., 2011)). Kefir grains (~0.1g) and pelleted fermented milk (from 1 ml sample) were suspended in 600 µl TES buffer (25mM Tris; 10mM EDTA; 50mM sucrose) with 25 units lyticase (Sigma-Aldrich, cat# L2524) and 20 mg/ml lysozyme (Sigma-Aldrich, cat# 62971) and incubated for 30 min at 37°C. The samples were then crushed with 0.3 g glass beads (Sigma-Aldrich, cat# G1277, 212-300 μm) at 4 m/s (fermented milk) and 6 m/s (kefir grains) for five times 20 seconds using a FastPrep®-24 instrument (MP Biomedicals). 150 µl 20 % SDS was added and after 5 min incubation at room temperature the tubes were centrifuged at maximum speed for 2 min. The supernatant was digested with 10 µl Proteinase K (20 mg/ml) for 30 min at 37°C and proteins were precipitated with 200 µl potassium acetate (5 M) for 15 min on ice. The samples were then centrifuged for 15 min at 4°C and the supernatant applied to phenol/chloroform extraction and DNA precipitation as described by (Kowalczyk et al., 2011). DNA quality was checked on agarose gel. DNA extraction from bacteria and yeasts was done as described above, using lysozyme and lyticase digestion, respectively.

**DNA extraction of isolated species** was performed using a simplified version of the extraction protocol used for kefir fermented milk containing removing the lysozyme digest (bacterial isolates) or the lyticase step (yeast isolates)

**Real-time PCR quantification (qPCR)** was done according to manufacturers instructions using the Applied Biosystems® StepOne™ Real-Time PCR System and the SYBR® Green RT-PCR Reagents Kit. The primer pairs LKF_KU504F - LKF_KU504R (Kim et al., 2015) and Kex-qPCR-forw (AGAGTAGTTCCTTTCACCCT TGCC) - Kex-qPCR-rev (AGTCGCTGGGTGATCGTCAG) were used for *L. kefiranofaciens* and *K. exigua*, respectively. Data was analyzed using the StepOne™ v2.3 software.

### Extracellular metabolite extraction

500 µl kefir fermented milk was centrifuged for 1 min at max. speed and 50 µl supernatant (kefir whey) were added to 950 µl 80°C H_2_O and incubated for 3 min at 80°C and 700 rpm. The samples were then centrifuged for 10 min at 0°C at maximum speed. The supernatant was harvested and ten times diluted for FIA-TOF measurement.

### FIA-qTOF MS untargeted measurements and data analysis

Metabolite extracts were injected into an Agilent 6550 time-of-flight mass spectrometer (ESI-iFunnel Q-TOF, Agilent Technologies) operated at 4 Ghz high resolution, in negative mode, with a mass / charge (*m/z*) range of 50−1,000 (Fuhrer et al., 2011). The mobile phase was 60:40 isopropanol:water (v/v) and 1 mM NH_4_F at pH 9.0 supplemented with hexakis(1H, 1H, 3H-tetrafluoropropoxy)phosphazine and 3-amino-1-propanesulfonic acid for online mass correction. Raw data was processed as described in (Fuhrer et al., 2011), and *m/z* features (ions) were annotated by matching their accurate mass-to-sum formulas of compounds to a custom KEGG database containing a merged database of *Escherichia coli* (eco), *Saccharomyces cerevisiae* (sce), and *Homo sapiens* (hsa) with 0.001 Da mass accuracy and accounting for deprotonation [M-H^+^]^−^. Only ions with high annotation accuracy were included in further analyses. This method has the inherent limitation that isobaric compounds, e.g., metabolites having identical *m/z* values, cannot be distinguished and that in-source fragmentation cannot be accounted for. The raw data of samples from the 2 sets of experiments (kefir development time course and *Lactococcus* – *Acetobacter* cross feeding) were processed and annotated separately to accommodate their different sample matrices or times of measurement. This raw data can be explored in Table S10. Raw data processing and annotation took place in MATLAB (MATLAB 2016b, The Mathworks, Natick) as described previously (Fuhrer et al., 2011), and downstream processing and statistical tests were performed in R (version 3.4.0, R Foundation for Statistical Computing, Vienna, Austria).

For the kefir development time course data ion intensities were transformed to Z-scores, filtered for non-changing patterns (Zimmermann et al., 2015) and the metabolite time sources were subjected to k-means clustering. In order to find the best K for clustering, we computed the Silhouette and Homogeneity score and performed the ensemble K-means clustering in varying random states to create a co-occurrence matrix (Figure S4). We chose six clusters because it returned the best compromise between the consistency and separability (Table S3). Overall dissimilarity of the ion profiles in different sampling points were visualized by Principal Component Analysis.

### qTRAP-MS targeted measurements

Metabolite extracts were injected on an Agilent HILIC Plus RRHD column (100 × 2.1mm × 1.8 μm; Agilent, Santa Clara, CA, USA). As was described previously, a gradient of mobile phase A (10 mM ammonium formate and 0.1% formic acid) and mobile phase B (acetonitrile with 0.1% formic acid) was used (Link et al., 2015). Metabolites were detected on a 5500 QTRAP triple quadrupole mass spectrometer in positive MRM scan mode (SCIEX, Framingham, MA, USA) at a constant flow rate of 400 μl/min.

### GC-MS measurements

Dried polar metabolites were derivatized with 50 μl of 20 mg/mL methoxyamine hydrochloride (Alfa Aesar, UK) solution in pyridine for 90 min at 37°C, followed by reaction with 100 μL N-methyl-trimethylsilyl-trifluoroacetamide (MSTFA) (Alfa Aesar, UK) for 10 hours at room temperature (Kanani and Klapa, 2007). GC-MS analysis was performed using a Shimadzu TQ8040 GC-(triple quadrupole) MS system (Shimadzu Corp.) equipped with a 30m × 0.25 mm × 0.25 µm DB-50MS capillary column (Phenomenex, USA). 1 μl of sample was injected in split mode (split ratio 1:20) at 250 °C using helium as a carrier gas with a flow rate of 1 ml/min. GC oven temperature was held at 100 °C for 4 min followed by an increase to 320 °C with a rate of 10 °C/min, and a final constant temperature period at 320 °C for 11 min. The interface and the ion source were held at 280 °C and 230 °C, respectively. The detector was operated both in scanning mode recording in the range of 50-600 m/z, as well as in MRM mode for specified metabolites. The metabolite quantification was carried out by calculating the area under the curve (AUC) of the identified marker ions of each metabolite normalized to the AUC of ribitol’s marker ion (marker ions m/z: γ-aminobutyric acid 174, aspartate 232, glycine 174, proline 142, ribitol 319). Data can be explored in table S13.

### Quantification of sugars and small organic acids by high-performance ion-chromatography

Samples were quenched with 4N sulfuric acid and deproteinated using an aqueous PCA solution. Arabinose and 2-hydroxyisobutyric acid were used as internal standards for the sugar and acid analysis, respectively. The diluted samples were analyzed on a Dionex ICS-3000 system (Thermo Fischer Scientific, Waltham (MA), USA). For the sugar analysis, an analytical anion-exchange column was used for the separation in combination with a pulsed amperometric detector (Corradini et al., 2012). The organic acids were analyzed in parallel on the same instrument, but with an analytical ion-exclusion column and a suppressed conductivity detector (Mullin and Emmons, 1997). Concentrations were calculated based on the chromatographic peak heights after normalizing to the internal standard (arabinose) using a one-point calibration. Raw data can be explored in tables S11 and S12.

### Amplicon analysis of 16S rRNA and ITS

All samples for the compositional analysis were examined by targeting the v4 regions of the 16S rRNA gene for bacteria and the ITS1/ITS2 region of the 18S rRNA gene for yeast. DNA was extracted as described above and cleaned with the DNA Clean & Concentrator™ kit (Zymo Research). Library preparation for the 16S V4 and 18S ITS was done using the NEXTflexTM 16S V4 Amplicon-Seq Kit 2.0 and NEXTflexTM 18S ITS Amplicon-Seq Kit, respectively. The barcoded amplicons were pooled in equal concentration and sequenced on the Illumina MiSeq platform using 2 × 250 bp at the EMBL genomic core facility in Heidelberg, Germany.

For the 16S analysis, raw paired-end sequences were quality-filtered using Cutadapt v1.10. (Martin, 2011) and merged using the Paired-End reAd mergeR (PEAR v0.9.6) (Zhang et al., 2014). QIIME2 (version 2018.04) was used for the downstream analysis (https://qiime2.org). The demultiplexed sequences were denoised and grouped into amplicon sequence variants (ASVs) using DADA2 (Callahan et al., 2016). The individual ASVs were taxonomically classified with 99% identity threshold by an open reference method (VSEARCH) using the 16S rRNA genes of the kefir isolates as reference (Rognes et al., 2016). Taxonomy of the non-kefir isolates ASVs were subsequently determined using the naive-Bayesian classifier trained on the Greengenes (Bokulich et al., 2018; McDonald et al., 2012). For the ITS analysis, raw sequences were quality-filtered using Cutadapt, but only ITS2-end reads were employed for the diversity analysis because the length variations in the ITS region resulted in the low number of the merged reads. As described above, the demultiplexed ITS2 sequences were denoised and subjected to taxonomic assignment by an open reference method, using the kefir isolated yeast as reference and UNITE for subsequent naive-Bayesian classifier (Kõljalg et al., 2013).

### Kefir and milk whey preparation

Kefir fermented milk was centrifuged for 30 min at 6000 rpm. The supernatant (whey) was filtered sterile with bottle top filters (pore size 0.22 µM). For milk whey preparation, 10 ml 32 % HCl was added to 1 liter UTH milk and stirred for ~10 min. The curdled milk was then centrifuged for 30 min at 6000 rpm and the pH of the supernatant was adjusted to 6.5 (pH of milk) with 10 M KOH. The whey was then centrifuged for another 30 min at 7000 rpm and filter sterilized. The sterile whey was stored at room temperature in the dark.

**Peptide-rich milk whey** was prepared by digestion of 1 liter UTH milk (3.5% fat) with 200 mg proteinase K for 72 hours. Digested milk was then centrifuged for 40 min at 7000 rpm and filter sterilized. The sterile whey was stored at room temperature in the dark.

### Species isolation from kefir grains and fermented milk

For species isolation, kefir fermented milk was diluted and plated directly, whereas kefir grains were homogenized using a Dounce tissue grinder before dilution. Diluted grain and milk fractions were inoculated in different rich media: GAM, mGAM, M17 supplemented with lactose and glucose, respectively, YPDA, MRS, TJA (tomato juice agar), YM (yeast mold agar), SD, Lactobacillus selective agar base (Neogen), and milk agar (200ml of 8% water agar plus 800ml UTH milk). MRS and TJA were used both, pure and diluted 1 to 1 with 48h kefir whey and 3.5% fat UTH milk, respectively. Growth was observed on plate for up to 10 days at room temperature, 30°C and 37°C, respectively. Kefir Lactobacilli were enriched using high salt conditions reaching from 1-5% NaCl.

Isolated species were identified by Sanger sequencing of the 16S/ITS region, using the primers S-D-Bact-0515-a-S-16 (GTGCCAGCMGCNGCGG) and S-*-Univ-1392-a-A-15 (ACGGGCGGTGTGTRC) (Klindworth et al., 2013). Unique isolates were sequenced using the Illumina® HiSeq 2000 platform.

### Pairwise species interactions in milk and on milk plate

The pairwise (binary) microbial interactions were assessed in liquid milk and on milk plates containing 0.04 g per liter bromocresol purple and blue, respectively. Binary species interactions in liquid milk were inoculated at final OD=0.025 each and grown at room temperature for 72 hours. Directly before DNA extraction, *E. coli* was spiked in at a final OD of 0.005 to allow relative quantification and growth assessment. DNA was extracted and the 16S amplicons were sequenced and their abundance quantified as described above. We removed samples with a small number of reads (<1,000), lacking *E. coli* sequences or contaminations above 5% total reads. The abundance of the inoculated species was converted into the fold-abundance of *E. coli* spike-in (the same amount in all samples). The normalized growth of the species in coculture was statistically compared to that in monoculture using t-test (p-value cutoff = 0.05). The effect of one species (A) on the other (B) was classified into positive, negative or neutral depending on growth change of the other species (ΔB). Interaction modes between two species were determined, on the basis of the reciprocal effects, to mutualism, competition, commensalism, amensalism, parasitism, and parabiosis (= neutralism) (Figure 4I).

Binary interactions on plates were assessed on milk agar medium containing 80 % UTH milk (3.5 % fat), 1 % agar and bromocresol blue as stain. The species to be tested were pinned on top of a background species spread on plate two hours prior to pinning. Pinning was done automated using the Singer ROTOR HDA Microbial Pinning Robot. Species were grown seven days at room temperature and plates were imaged under controlled lighting conditions (spImager S&P Robotics Inc.) using an 18 megapixel Canon Rebel T3i. Colony surface area was determined semi-automated using ImageJ. The growth effects (background species to pinned species) were termed positive, negative, or neutral, respectively, if the colony surface area of pinned species was significantly larger, smaller, or similar on plates with background species compared to plates without background species (t-test with p-value cutoff of 0.05). Interaction modes were coined as described above.

### Metabolic binary species interactions in milk

The 20 most interesting species (most abundant and isolated from our reference kefir GER6, respectively) involved in kefir fermentation, including several rare species, were inoculated in milk stained with bromocresol purple solution, a pH indicator, that is yellow at a pH below 5.2 and purple above pH 6.8. The inoculated species were grown for 76 hours at room temperature. Microbial growth was assessed by pH-dependent color change and monitored by automated hourly scanning with a standard flatbed scanner and subsequent analysis of pH changes using the growth profiler software (Enzyscreen B.V., Heemstede, Netherlands). The species were either inoculated alone or in pairs and the potential of the pairs to acidify milk within 76 hours was compared with single inoculate.

The RGB code for changing color in response to different pH were independently plotted, showing that the R-values are linearly-correlated with pH change in the range of 3 to 7. Thus, the acidification by inoculated species was measured by converting the R-value to pH. The species were either inoculated alone or in pairs. The experiment was performed three times, with each species tested in duplicate per run. The interaction potential of the pairs for milk acidification was determined on the basis of reduction area under the pH curves within 76 hours. We define the positive interaction if the acidification area of coculture is bigger than the sum of those of single inoculum and the negative interaction if the pH of coculture is higher than those of both individuals.

***L. lactis* spent whey** was produced by *L. lactis* (strains SB-17, SB-261 and SB-352) fermentation for 72 hours or by *L. lactis* (strain SB-150) for 120 hours at 30°C. Whey was harvested by centrifugation for 40 min at 7000 rpm and filter sterilized. The sterile whey was stored at room temperature in the dark.

### Metabolomics measurements in milk and *L. lactis* spent whey

Four *Acetobacter* isolates (*A. fabarum*: internal stock IDs SB-290, SB-373 and *A. ghanensis*: internal stock IDs SB-354, SB-380) were grown in whey produced with four *L. lactis* isolates, while sampling extracellular metabolomics samples over time. These samples were measured with untargeted metabolomics in an independent metabolomics experiment. Possibly exchanged metabolites were identified by selecting ions that showed an increase during *L. lactis* growth (log2(FC) ≥ 1) and a decrease during *Acetobacter* growth (log2(FC) ≤ 1).

### Estimation of amino acid uptake and secretion rates

We built genome-scale metabolic models for all bacterial species (which account for more than 90% of the total community composition) using CarveMe (Machado et al., 2018). The models were used to simulate the expected uptake/secretion rates of amino acids using flux balance analysis. At each time point the individual rates for each species were combined with the measured species abundances to estimate the overall net accumulation/depletion rates for all amino acids.

### Estimation of expected amino acid accumulation

We estimated the level of accumulation of amino acids that would be expected by proteolysis alone. For this we normalized the measured accumulation rates by the natural amino acid abundance in milk protein and estimated a first-order kinetic rate for proteolysis (1.36e-4 h-1) by least-squares regression. We then multiplied this value by the natural abundance of each amino acid to estimate their respective accumulation rate by proteolysis.

