## Supplementary Figures S1 - S24 for "Emergence of stable coexistence in a complex microbial community through metabolic cooperation and spatio-temporal niche partitioning"

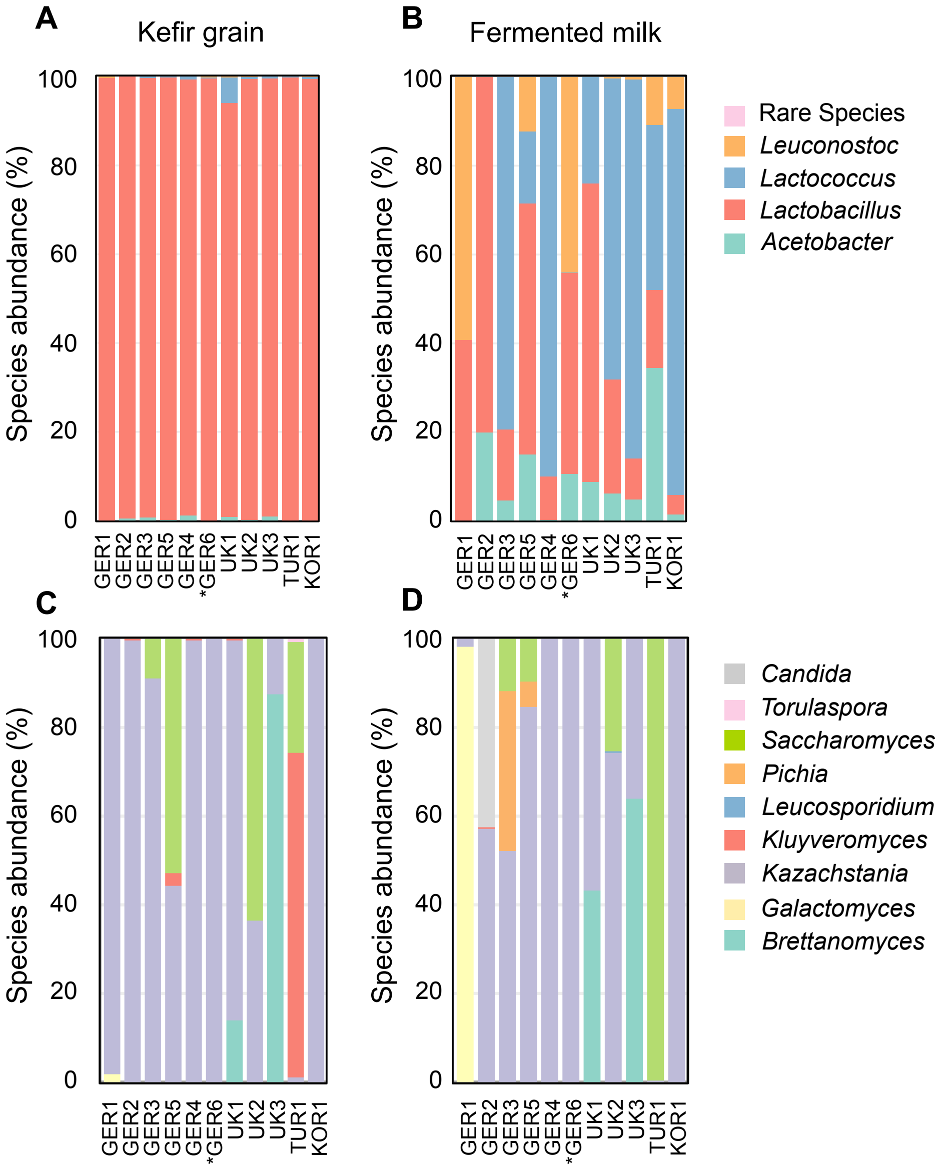


**Figure S1.** Microbial composition of the kefir grain (A,C) and kefir fermented milk (B,D) obtained from diverse countries. Bacterial (A, B) and fungal (C, D) populations were measured using 16S rDNA and ITS amplicons. Rare species = relative abundance <0.01; GER6* = reference kefir grain (GER6, OG2) used for the kefir fermentation curve.

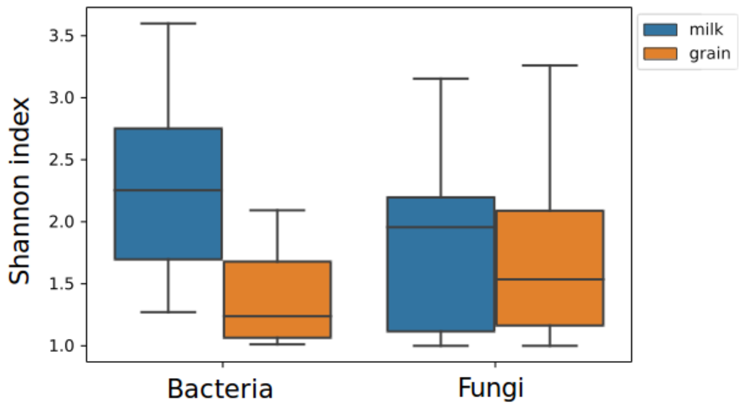


**Figure S2.** Variation in the bacterial and fungal Shannon diversity index of the kefir grain and kefir fermented milk.

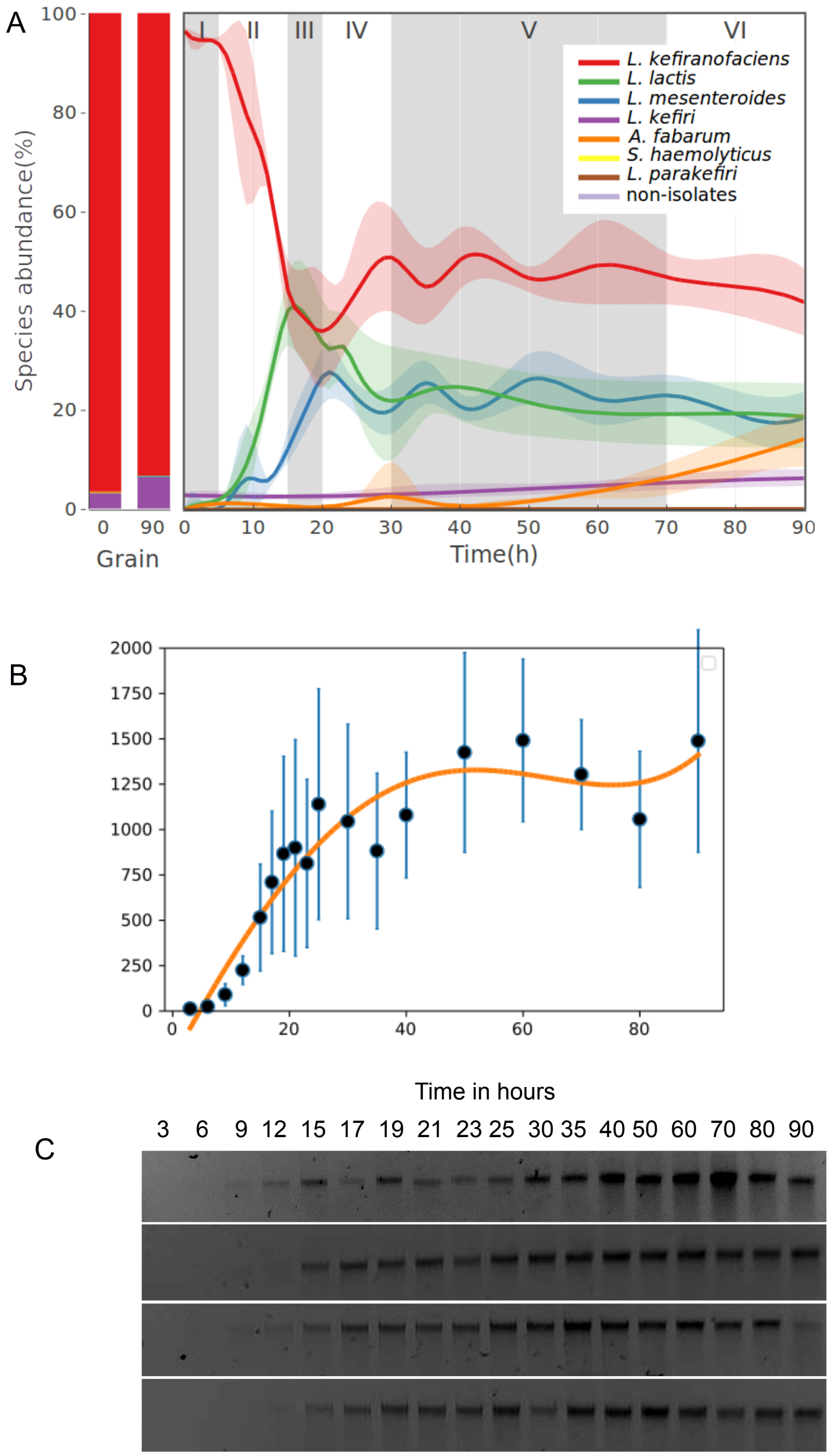


**Figure S3.** Additional information related to Figure 1D. (A) Temporal dynamics of bacterial composition during fermentation in the kefir fermented milk assessed by 16S amplicon sequencing. The non-isolates represent the sum of all species, which are not in the kefir isolate collection. The fermentation is split into 6 different phases depending on the abundance changes of the major species. (B) Fitting of DNA concentrations to sigmoid curve, (C) DNA extracted from fermentation samples. DNA concentration was used to quantify kefir species.

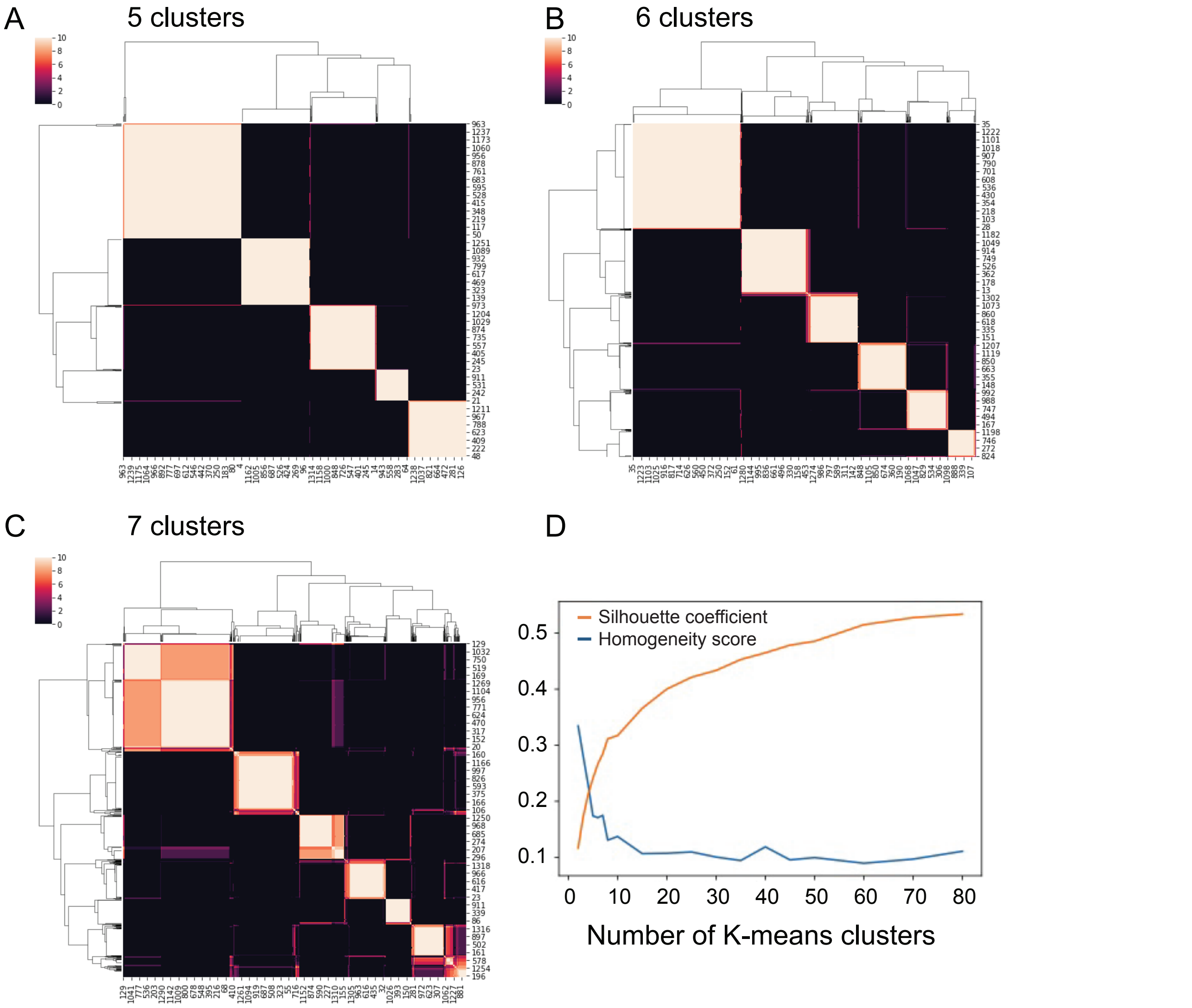


**Figure S4.** K-means clustering of untargeted metabolomics data in (A) 5, (B) 6 and (C) 7 different clusters. Splitting in six different clusters was found to be the highest number that still allowed clean separation of metabolites into distinct clusters. (D) The graph shows the change in silhouette coefficient and homogeneity score as the number of clusters is increased.

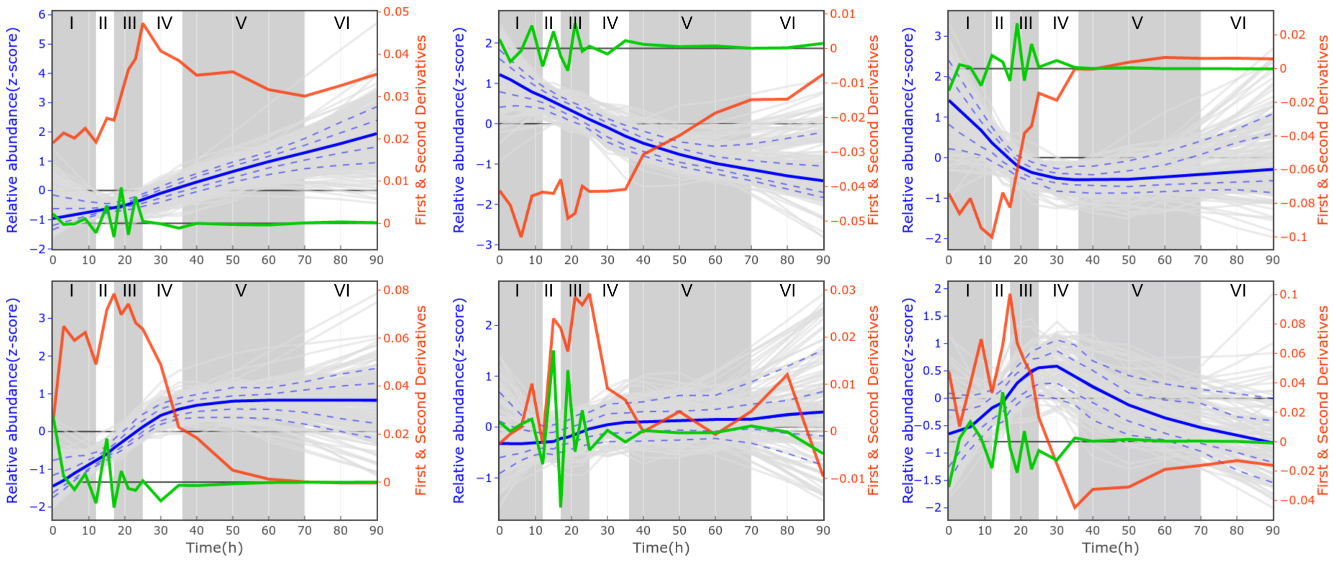


**Figure S5.** First and second derivatives of metabolite dynamics match kefir fermentation phases.

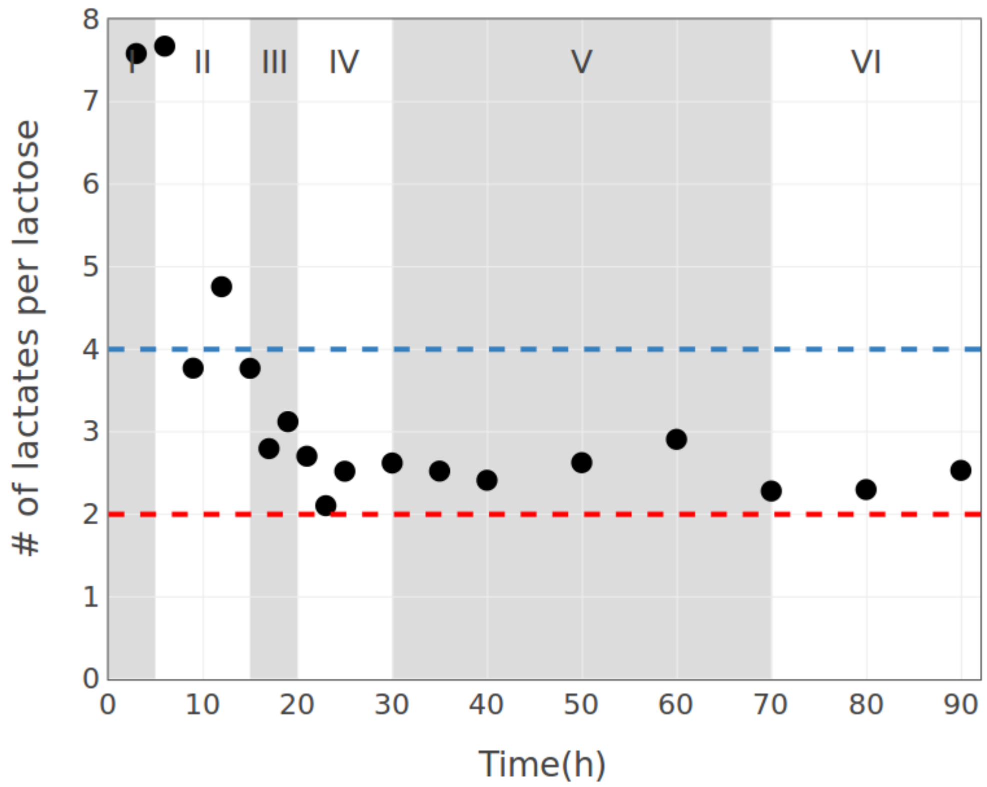


**Figure S6.** Conversion of lactose into lactate during kefir fermentation. The blue and the red lines show the production of four and two lactates per lactose molecule, as typical for homofermentative and heterofermentative LAB, respectively. Note that the higher yields at initial time points are due to lactate from inoculum and likely measurement errors at very low concentrations.

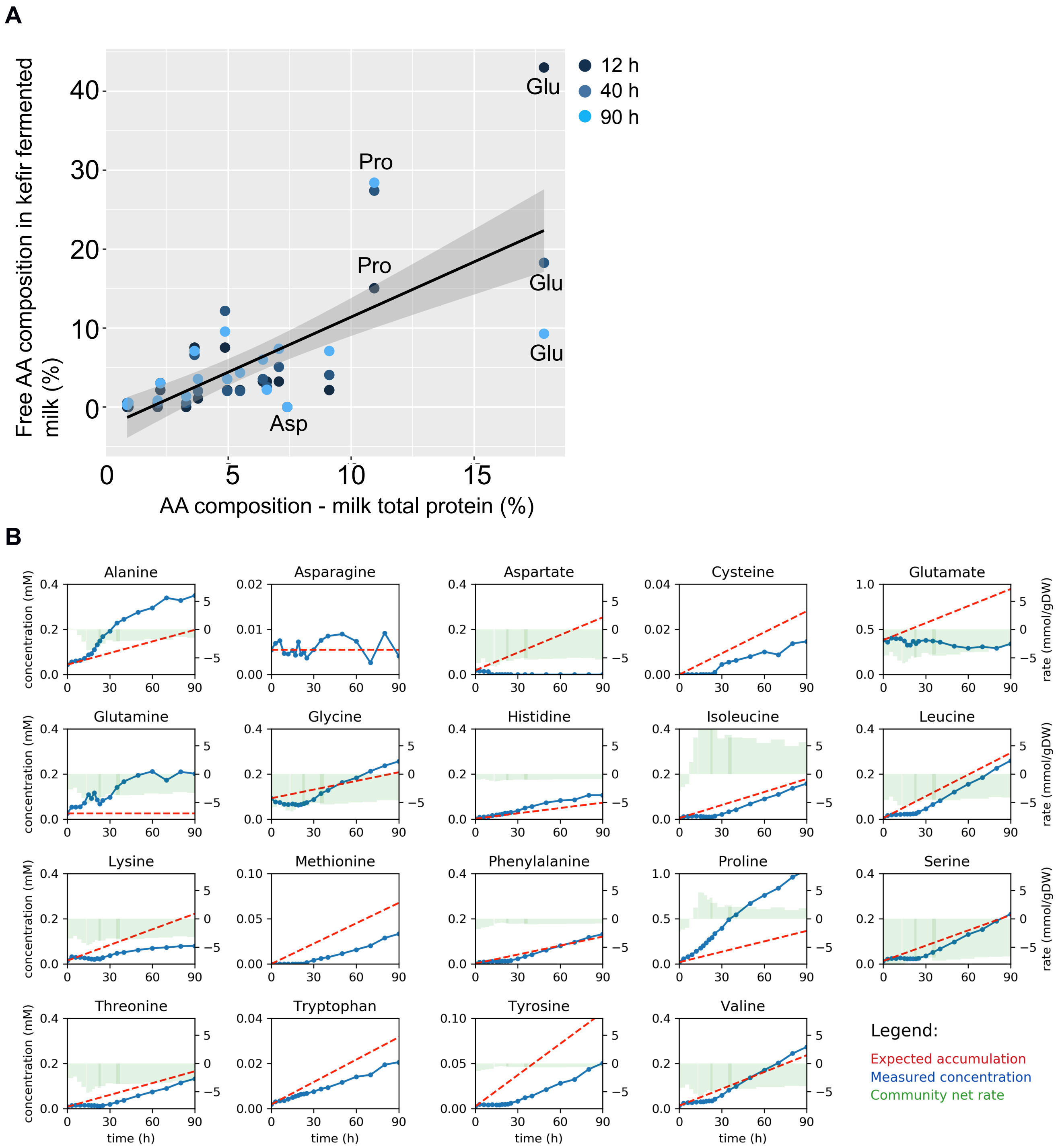


**Figure S7.** (A) Amino acid composition of hydrolyzed milk total protein (%) versus composition of free amino acids in kefir (%) released after 12, 40 and 90 hours fermentation.
Milk amino acid composition was averaged from (Park, 2007; Schönfeldt et al., 2011).
(B) Comparison of expected accumulation (red dotted lines) and measured concentrations (blue lines) of amino acids in milk kefir. Green bars indicate the model-based estimation of uptake (negative values) and secretion rates (positive values).

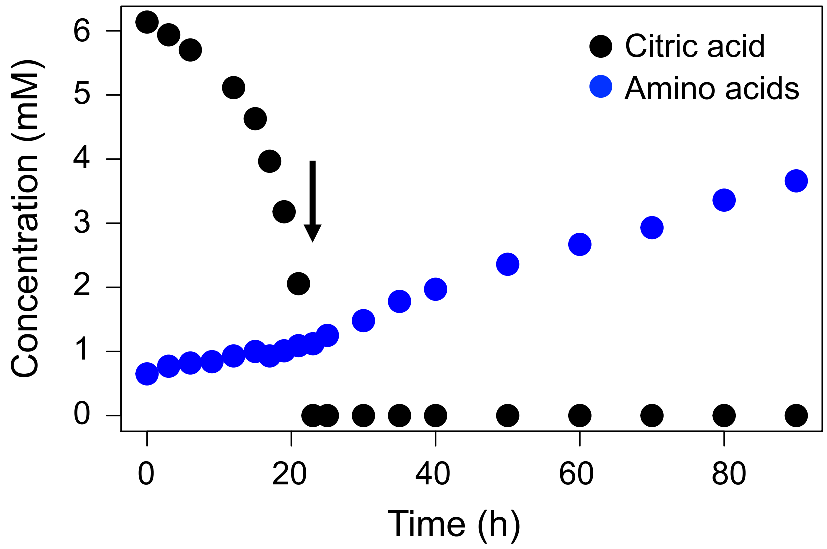


**Figure S8.** Increase in free amino acids after citrate depletion during kefir fermentation. Amino acids = total amount of free amino acids in kefir fermented milk.

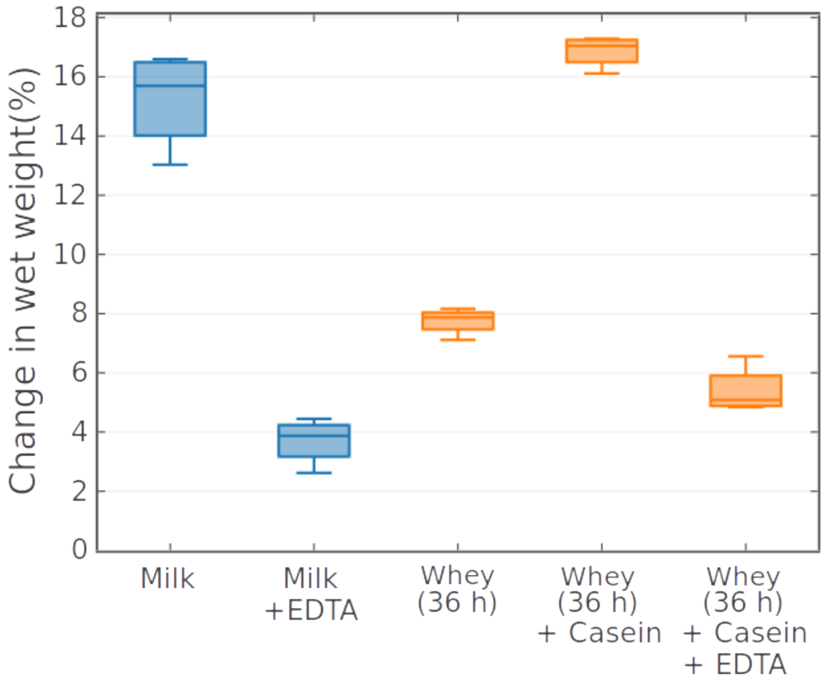


**Figure S9.** Effect of EDTA and protein on grain wet weight gain after 72 hours fermentation. Kefir grains grown in whey harvested after 36 hours fermentation reveal decreased growth that is restored by casein supplementation. Addition of EDTA inhibits grain growth in both, milk and casein-supplemented kefir whey.

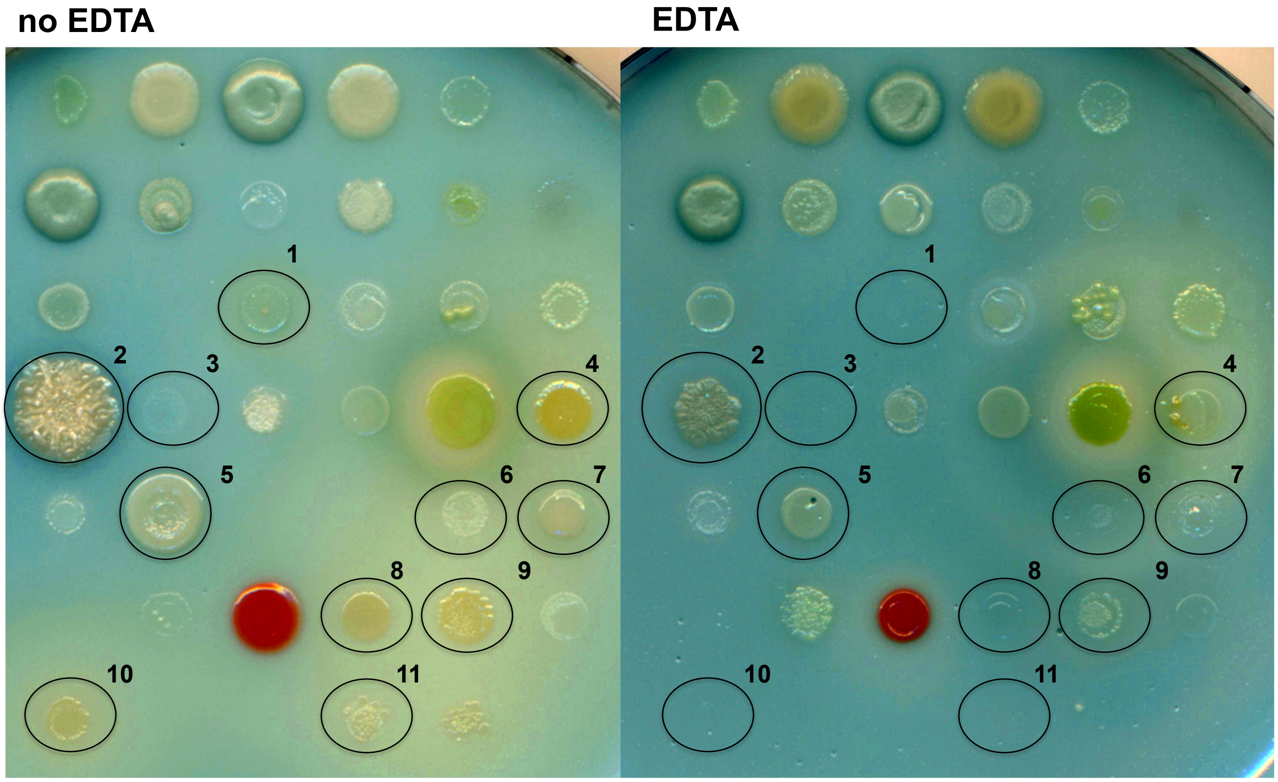


| | | | |
|---|---|---|---|
| 1 | *Leuconostoc mesenteroides* (SB-230a) | 7 | *Micrococcus luteus* (SB-332) |
| 2 | *Bacillus simplex* (SB-271) | 8 | *Acetobacter ghanensis* (SB-354) |
| 3 | *Lactobacillus parakefiri* (SB-280) | 9 | *Acetobacter fabarum* (SB-373) |
| 4 | *Acetobacter fabarum* (SB-290) | 10 | *Acetobacter ghanensis* (SB-380) |
| 5 | *Brevibacterium casei* (SB-303) | 11 | *Moraxella osloensis* (SB-424) |
| 6 | *Lactobacillus parakefiri* (SB-325) | | |

**Figure S10.** Inhibition of kefir species by EDTA on milk-agar colored with bromocresol green. Most affected species are circled and listed below.

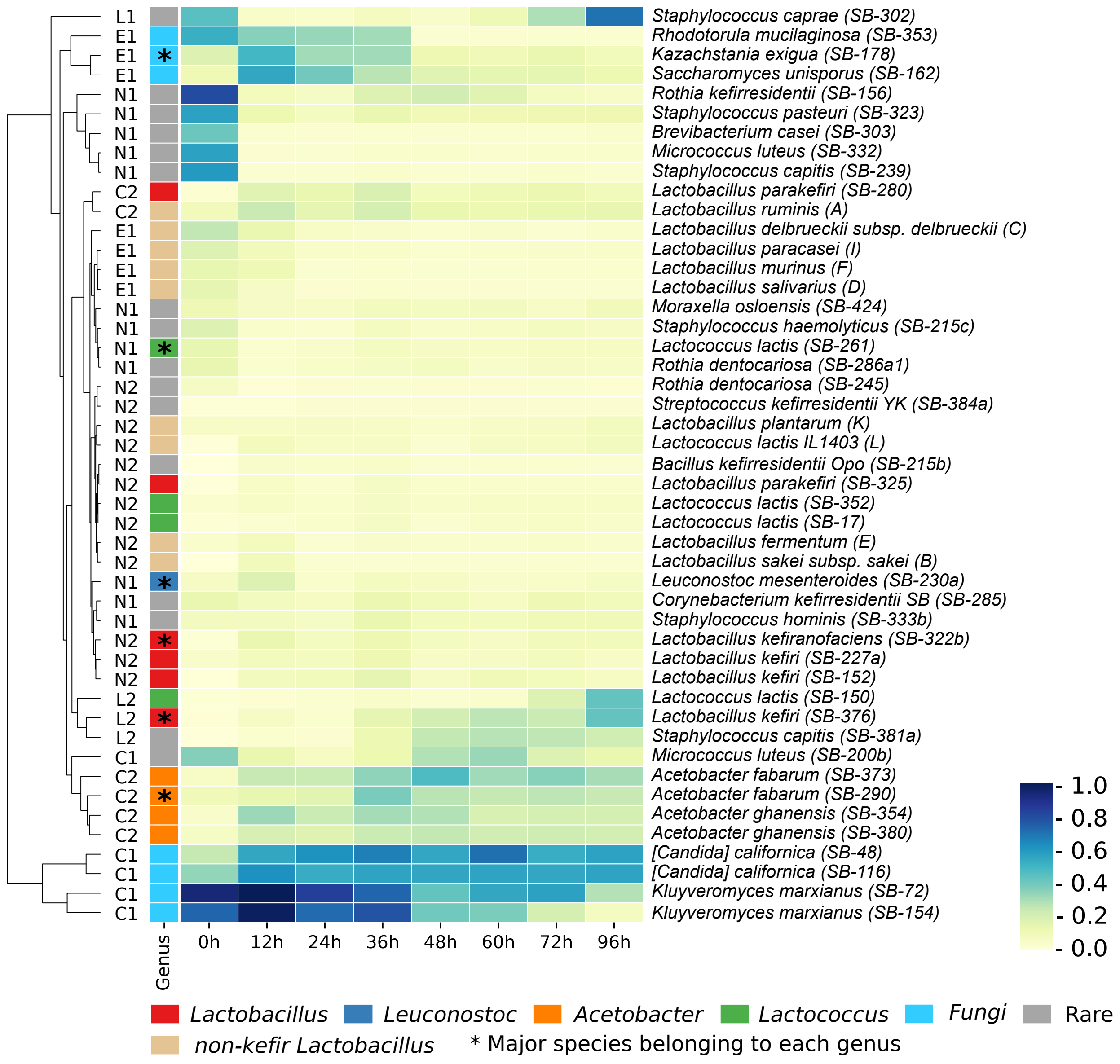


**Figure S11.** Growth (OD) of individual isolates in kefir spent whey harvested at different fermenting times. "0h" refers to non-fermented milk whey at pH 6.5. Groups: E=early, L=late, C=constant, N= non- or decreased growth, subgroup: 1=growth in milk whey, not in kefir whey, 2=no growth in milk whey.

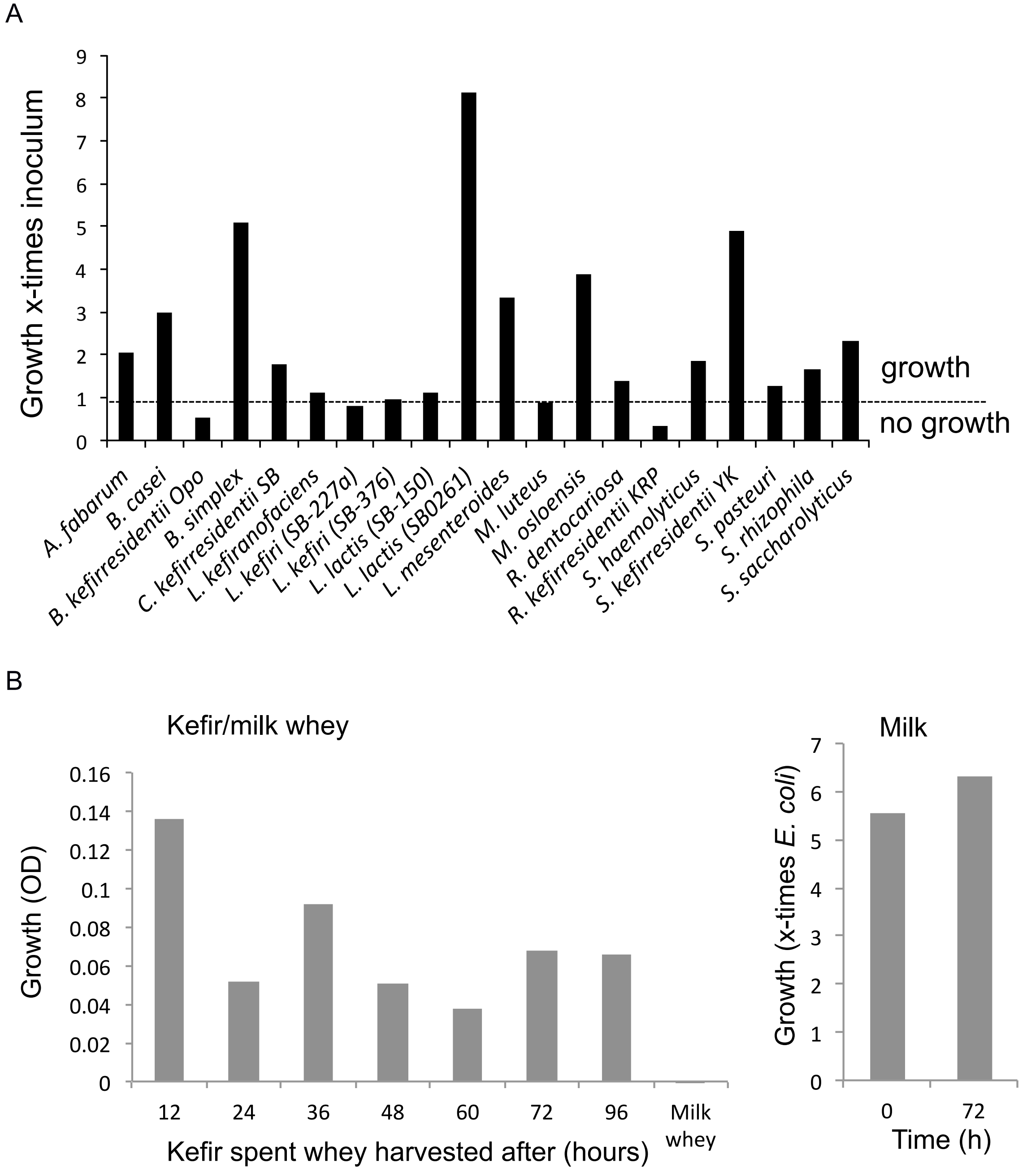


**Figure S12.** Growth of kefir species (A) in milk determined by 16S amplicon sequencing and *E. coli* standard. (B) Growth of *L. kefiranofaciens* in kefir spent whey, milk whey (pH= 6.5) (after 7 days) and in milk after 72 hours.

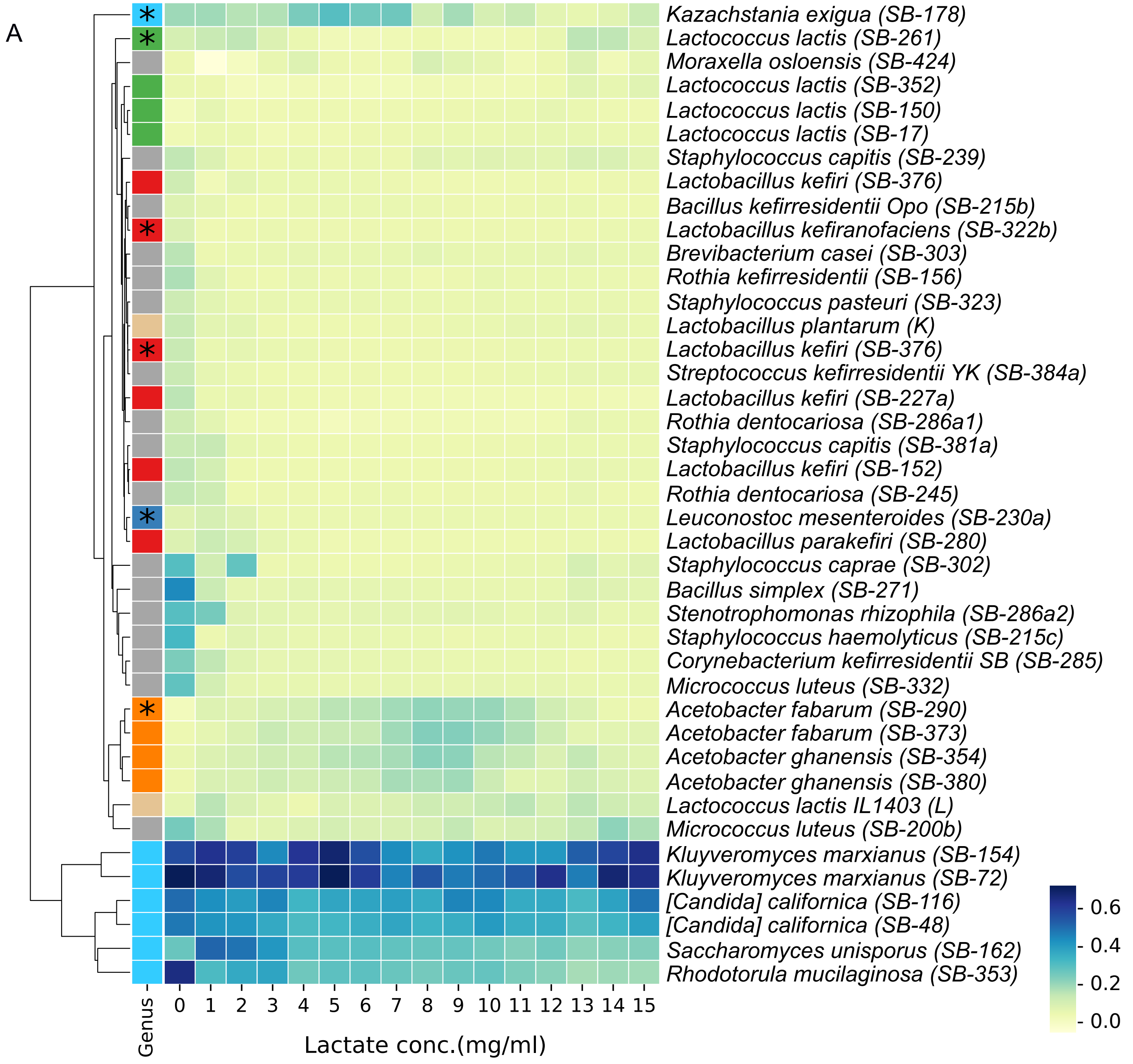
A
Kazachstania exigua (SB-178)
Lactococcus lactis (SB-261)
Moraxella osloensis (SB-424)
Lactococcus lactis (SB-352)
Lactococcus lactis (SB-150)
Lactococcus lactis (SB-17)
Staphylococcus capitis (SB-239)
Lactobacillus kefiri (SB-376)
Bacillus kefirresidentii Opo (SB-215b)
Lactobacillus kefiranofaciens (SB-322b)
Brevibacterium casei (SB-303)
Rothia kefirresidentii (SB-156)
Staphylococcus pasteuri (SB-323)
Lactobacillus plantarum (K)
Lactobacillus kefiri (SB-376)
Streptococcus kefirresidentii YK (SB-384a)
Lactobacillus kefiri (SB-227a)
Rothia dentocariosa (SB-286a1)
Staphylococcus capitis (SB-381a)
Lactobacillus kefiri (SB-152)
Rothia dentocariosa (SB-245)
Leuconostoc mesenteroides (SB-230a)
Lactobacillus parakefiri (SB-280)
Staphylococcus caprae (SB-302)
Bacillus simplex (SB-271)
Stenotrophomonas rhizophila (SB-286a2)
Staphylococcus haemolyticus (SB-215c)
Corynebacterium kefirresidentii SB (SB-285)
Micrococcus luteus (SB-332)
Acetobacter fabarum (SB-290)
Acetobacter fabarum (SB-373)
Acetobacter ghanensis (SB-354)
Acetobacter ghanensis (SB-380)
Lactococcus lactis IL1403 (L)
Micrococcus luteus (SB-200b)
Kluyveromyces marxianus (SB-154)
Kluyveromyces marxianus (SB-72)
[Candida] californica (SB-116)
[Candida] californica (SB-48)
Saccharomyces unisporus (SB-162)
Rhodotorula mucilaginosa (SB-353)
Genus
0 1 2 3 4 5 6 7 8 9 10 11 12 13 14 15
Lactate conc.(mg/ml)
0.6
0.4
0.2
0.0

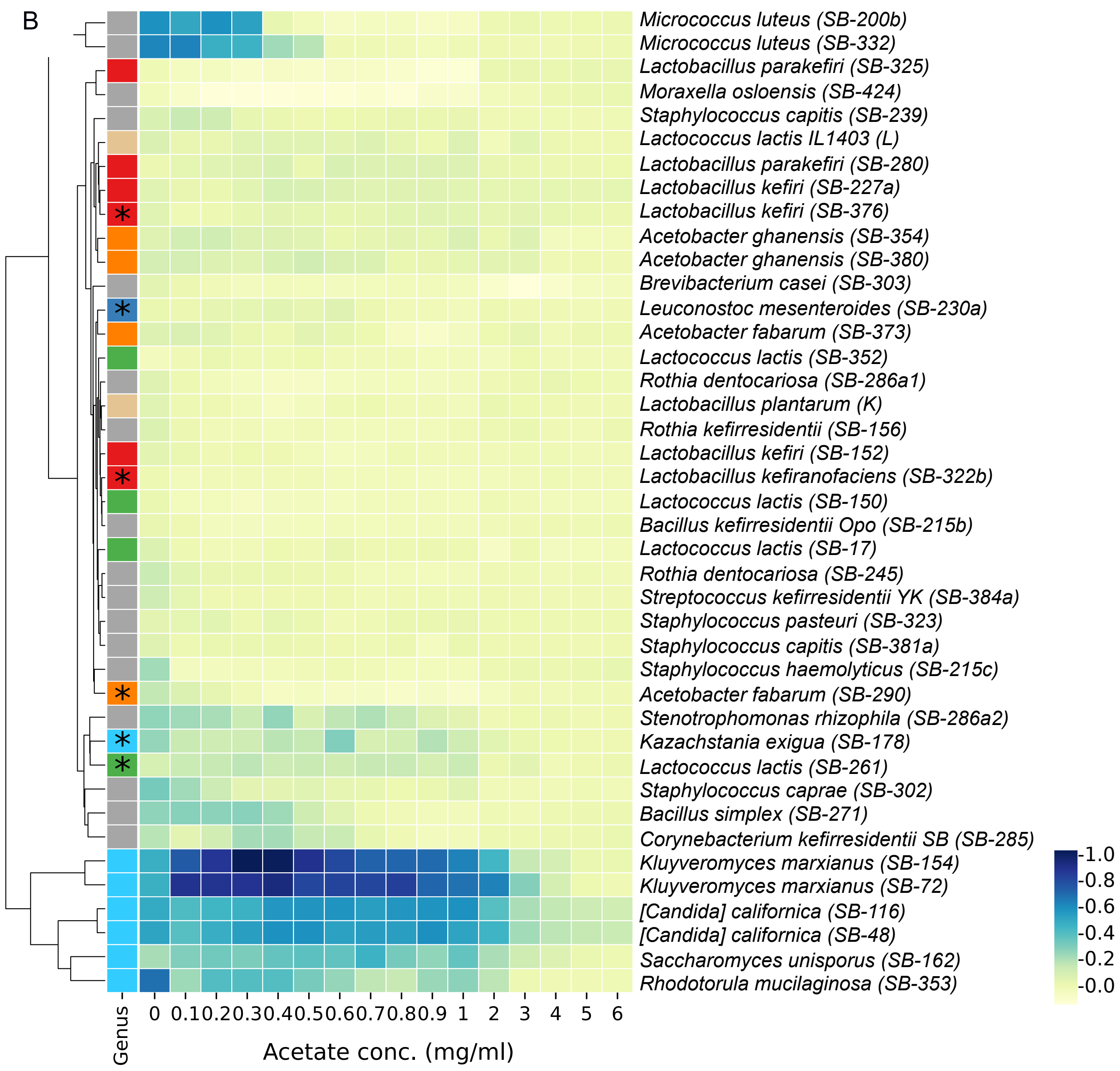
B
Micrococcus luteus (SB-200b)
Micrococcus luteus (SB-332)
Lactobacillus parakefiri (SB-325)
Moraxella osloensis (SB-424)
Staphylococcus capitis (SB-239)
Lactococcus lactis IL1403 (L)
Lactobacillus parakefiri (SB-280)
Lactobacillus kefiri (SB-227a)
Lactobacillus kefiri (SB-376)
Acetobacter ghanensis (SB-354)
Acetobacter ghanensis (SB-380)
Brevibacterium casei (SB-303)
Leuconostoc mesenteroides (SB-230a)
Acetobacter fabarum (SB-373)
Lactococcus lactis (SB-352)
Rothia dentocariosa (SB-286a1)
Lactobacillus plantarum (K)
Rothia kefirresidentii (SB-156)
Lactobacillus kefiri (SB-152)
Lactobacillus kefiranofaciens (SB-322b)
Lactococcus lactis (SB-150)
Bacillus kefirresidentii Opo (SB-215b)
Lactococcus lactis (SB-17)
Rothia dentocariosa (SB-245)
Streptococcus kefirresidentii YK (SB-384a)
Staphylococcus pasteuri (SB-323)
Staphylococcus capitis (SB-381a)
Staphylococcus haemolyticus (SB-215c)
Acetobacter fabarum (SB-290)
Stenotrophomonas rhizophila (SB-286a2)
Kazachstania exigua (SB-178)
Lactococcus lactis (SB-261)
Staphylococcus caprae (SB-302)
Bacillus simplex (SB-271)
Corynebacterium kefirresidentii SB (SB-285)
Kluyveromyces marxianus (SB-154)
Kluyveromyces marxianus (SB-72)
[Candida] californica (SB-116)
[Candida] californica (SB-48)
Saccharomyces unisporus (SB-162)
Rhodotorula mucilaginosa (SB-353)
1.0
0.8
0.6
0.4
0.2
0.0
Genus
0 0.1 0.2 0.3 0.4 0.5 0.6 0.7 0.8 0.9 1 2 3 4 5 6
Acetate conc. (mg/ml)

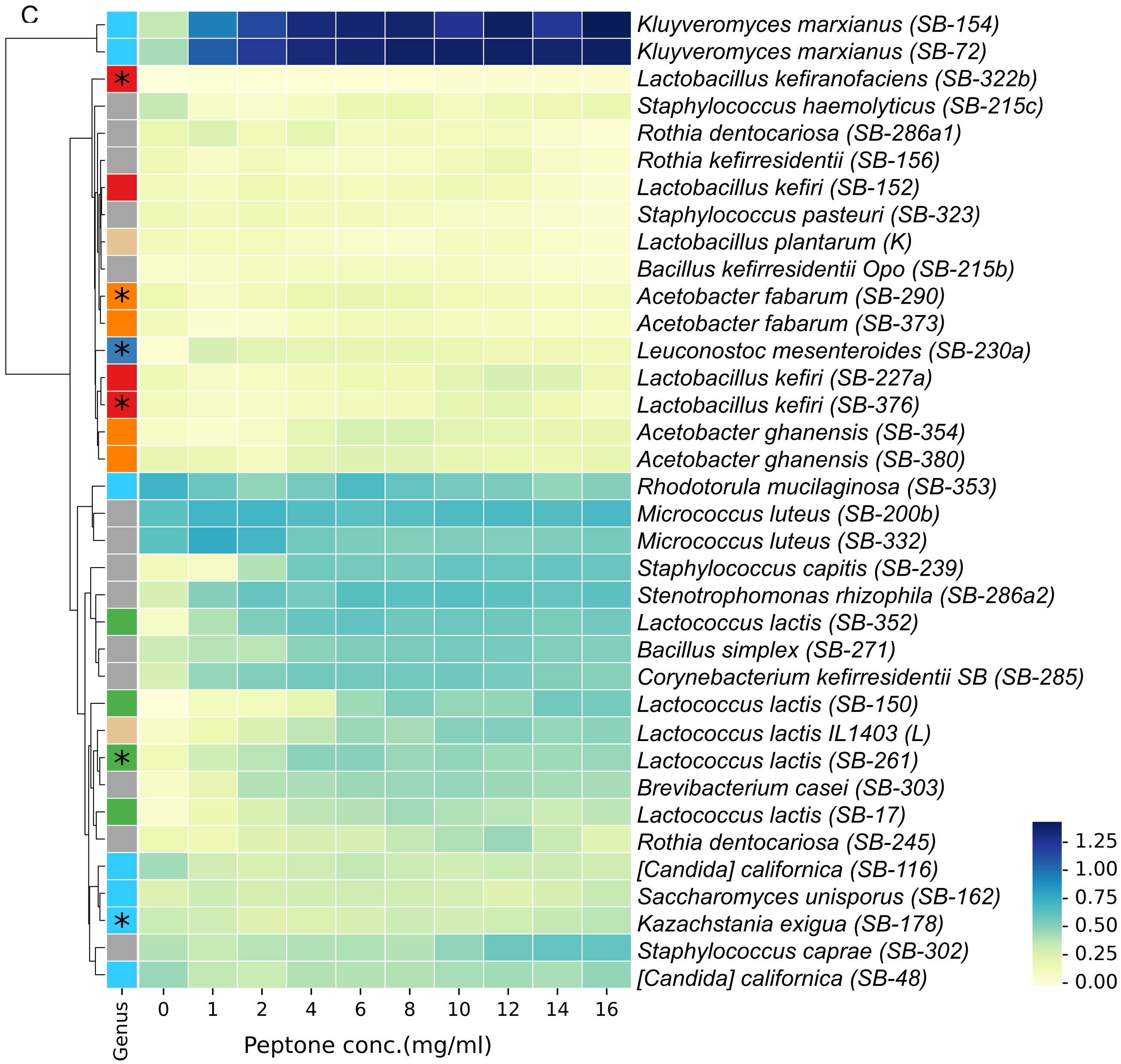
C
Kluyveromyces marxianus (SB-154)
Kluyveromyces marxianus (SB-72)
Lactobacillus kefiranofaciens (SB-322b)
Staphylococcus haemolyticus (SB-215c)
Rothia dentocariosa (SB-286a1)
Rothia kefirresidentii (SB-156)
Lactobacillus kefiri (SB-152)
Staphylococcus pasteuri (SB-323)
Lactobacillus plantarum (K)
Bacillus kefirresidentii Opo (SB-215b)
Acetobacter fabarum (SB-290)
Acetobacter fabarum (SB-373)
Leuconostoc mesenteroides (SB-230a)
Lactobacillus kefiri (SB-227a)
Lactobacillus kefiri (SB-376)
Acetobacter ghanensis (SB-354)
Acetobacter ghanensis (SB-380)
Rhodotorula mucilaginosa (SB-353)
Micrococcus luteus (SB-200b)
Micrococcus luteus (SB-332)
Staphylococcus capitis (SB-239)
Stenotrophomonas rhizophila (SB-286a2)
Lactococcus lactis (SB-352)
Bacillus simplex (SB-271)
Corynebacterium kefirresidentii SB (SB-285)
Lactococcus lactis (SB-150)
Lactococcus lactis IL1403 (L)
Lactococcus lactis (SB-261)
Brevibacterium casei (SB-303)
Lactococcus lactis (SB-17)
Rothia dentocariosa (SB-245)
[Candida] californica (SB-116)
Saccharomyces unisporus (SB-162)
Kazachstania exigua (SB-178)
Staphylococcus caprae (SB-302)
[Candida] californica (SB-48)
1.25
1.00
0.75
0.50
0.25
0.00
Genus
0
1
2
4
6
8
10
12
14
16
Peptone conc.(mg/ml)

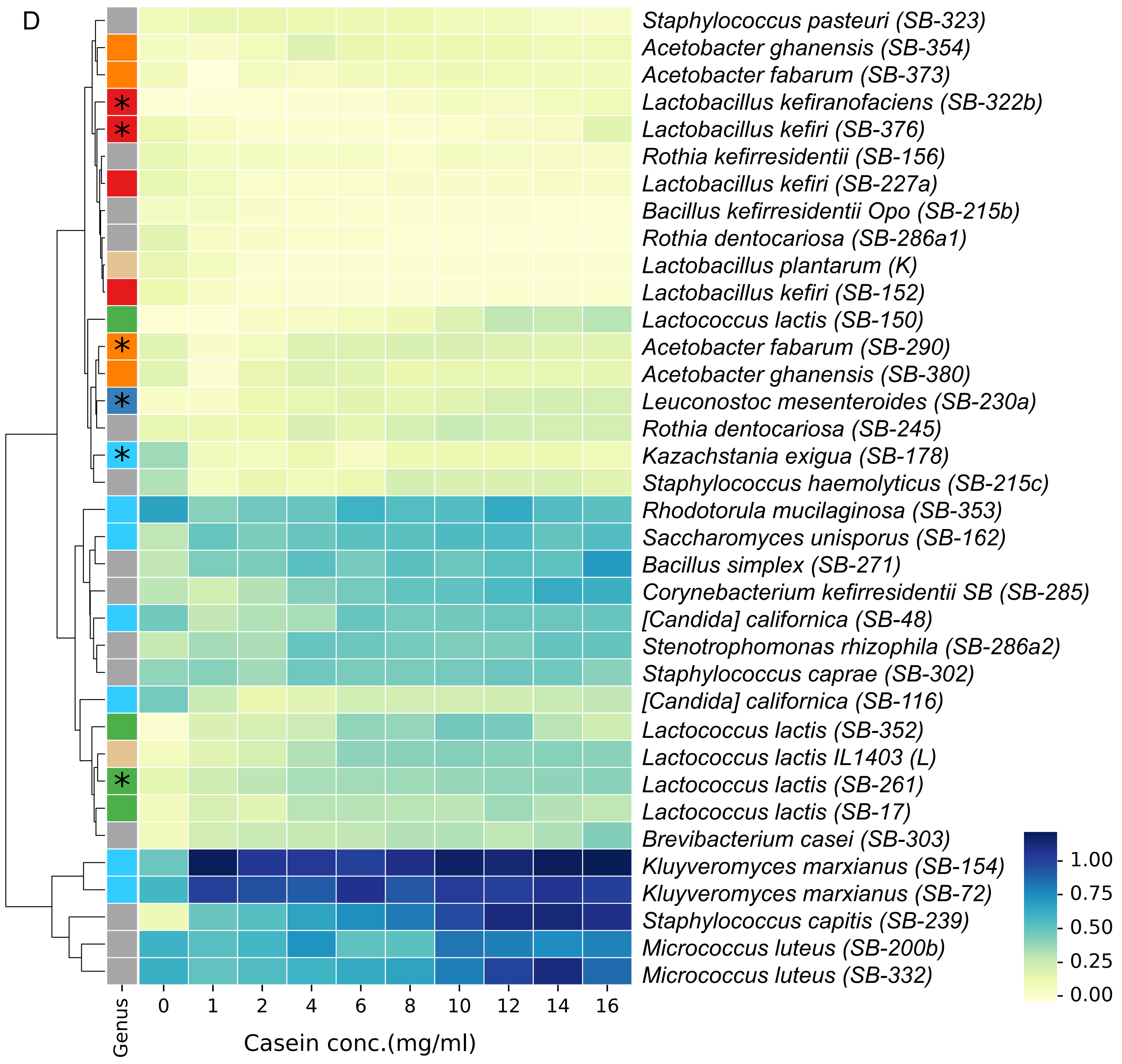
D
Staphylococcus pasteuri (SB-323)
Acetobacter ghanensis (SB-354)
Acetobacter fabarum (SB-373)
Lactobacillus kefiranofaciens (SB-322b)
Lactobacillus kefiri (SB-376)
Rothia kefirresidentii (SB-156)
Lactobacillus kefiri (SB-227a)
Bacillus kefirresidentii Opo (SB-215b)
Rothia dentocariosa (SB-286a1)
Lactobacillus plantarum (K)
Lactobacillus kefiri (SB-152)
Lactococcus lactis (SB-150)
Acetobacter fabarum (SB-290)
Acetobacter ghanensis (SB-380)
Leuconostoc mesenteroides (SB-230a)
Rothia dentocariosa (SB-245)
Kazachstania exigua (SB-178)
Staphylococcus haemolyticus (SB-215c)
Rhodotorula mucilaginosa (SB-353)
Saccharomyces unisporus (SB-162)
Bacillus simplex (SB-271)
Corynebacterium kefirresidentii SB (SB-285)
[Candida] californica (SB-48)
Stenotrophomonas rhizophila (SB-286a2)
Staphylococcus caprae (SB-302)
[Candida] californica (SB-116)
Lactococcus lactis (SB-352)
Lactococcus lactis IL1403 (L)
Lactococcus lactis (SB-261)
Lactococcus lactis (SB-17)
Brevibacterium casei (SB-303)
Kluyveromyces marxianus (SB-154)
Kluyveromyces marxianus (SB-72)
Staphylococcus capitis (SB-239)
Micrococcus luteus (SB-200b)
Micrococcus luteus (SB-332)
1.00
0.75
0.50
0.25
0.00
Genus
0
1
2
4
6
8
10
12
14
16
Casein conc.(mg/ml)

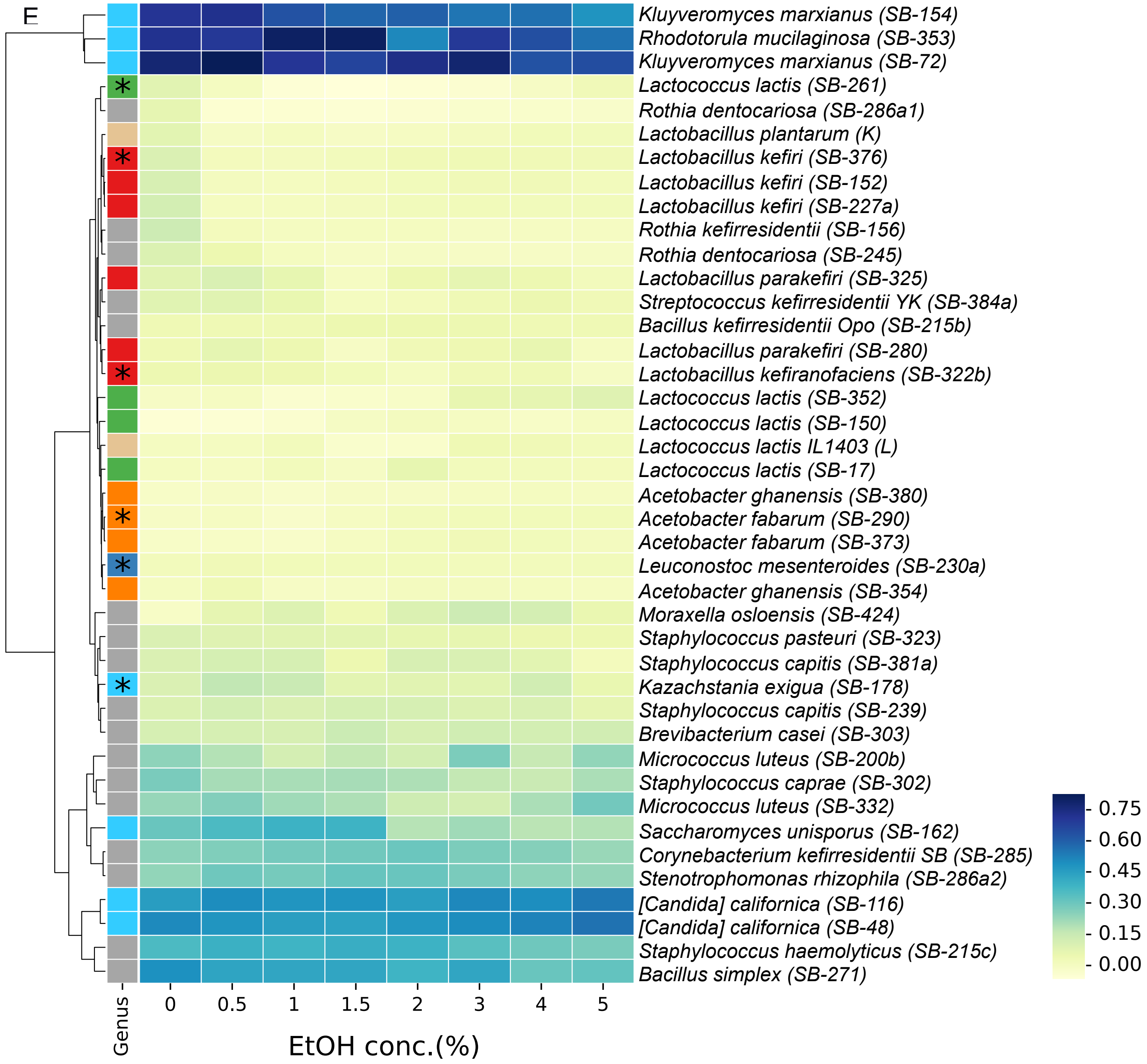


**Figure S13.** Growth of different species in substrate-amended milk whey. (A) Lactate (B) Acetate (C) Peptone (D) Casein and (E) Ethanol supplementation.

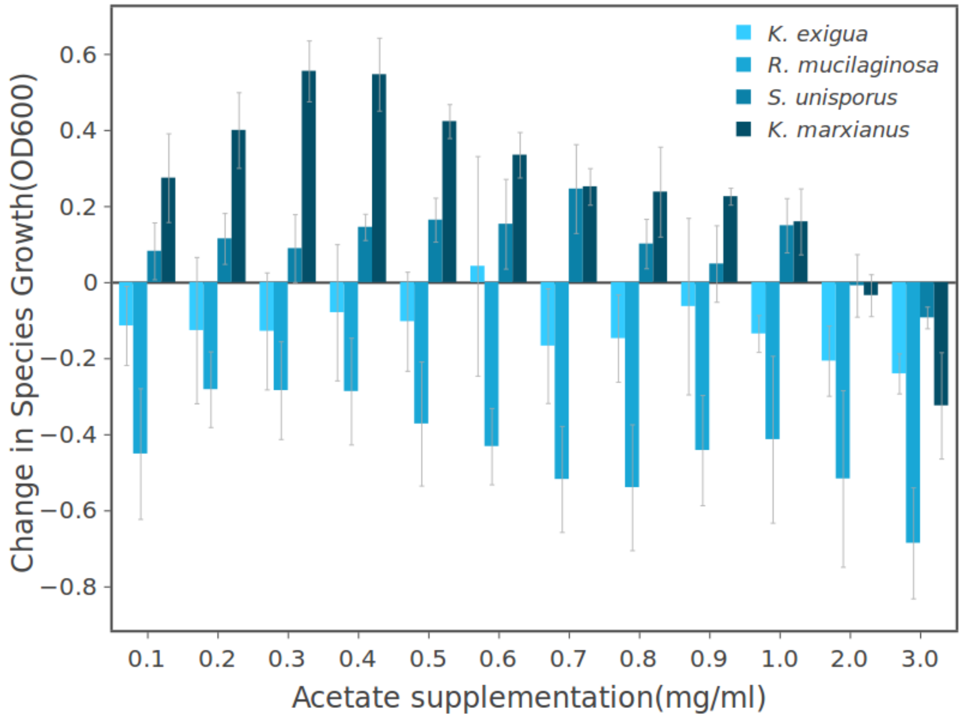


**Figure S14.** Effect of acetate on growth of *K. exigua*, *R. mucilaginosa*, *S. unisporus* and *K. marxianus*. *S. unisporus* and *K. marxianus* profit from low acetate concentrations, *K. exigua* and *R. mucilaginosa*, are inhibited even by small acetate supplements. Changes in species growth are assessed relative to growth in non-supplemented MW.

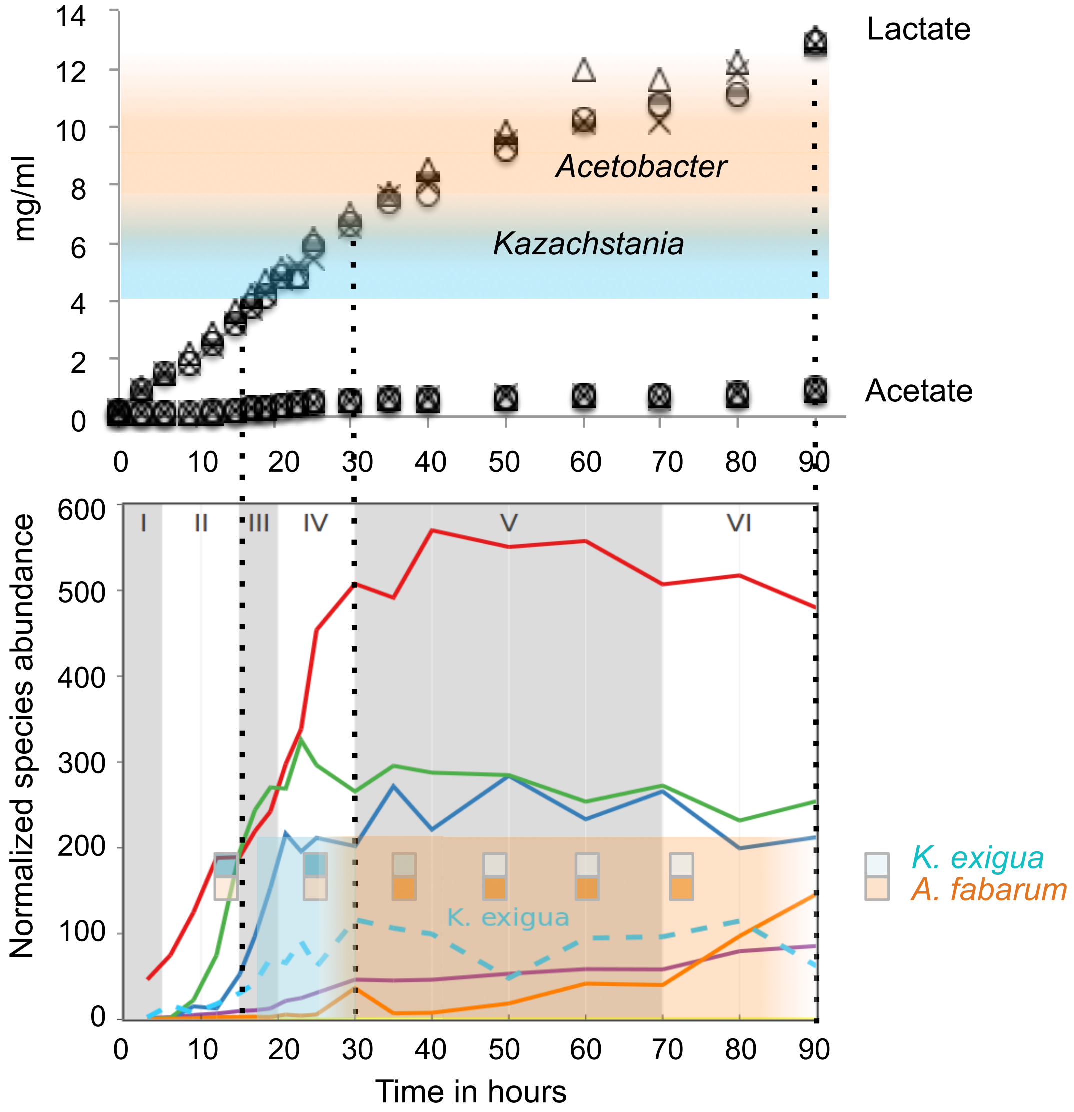


**Figure S15**. Lactate shapes consecutive growth windows of *K. exigua* (yeast) and *A. fabarum.* Overlay of lactate and acetate concentration during kefir fermentation with species preferred growth time and its optimal supportive amount of lactate in milk whey feeding assay. The blue and orange bars represent the growth of *K. exigua* and *A. fabarum* in kefir spent whey at corresponding time points, darker colors represent increased growth (OD). Full dataset can be found in Figure S13.

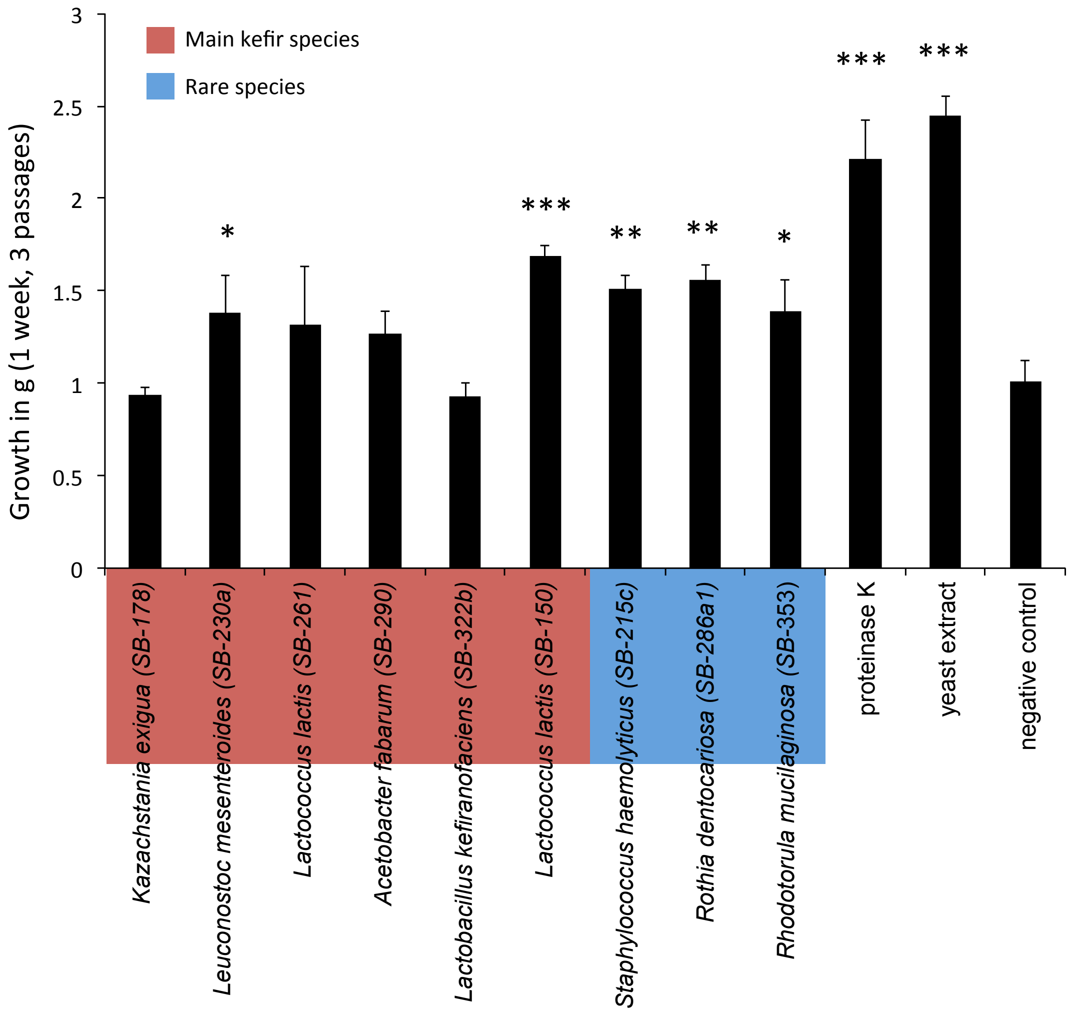


**Figure S16.** Kefir grain growth profit from rare species and supplements.

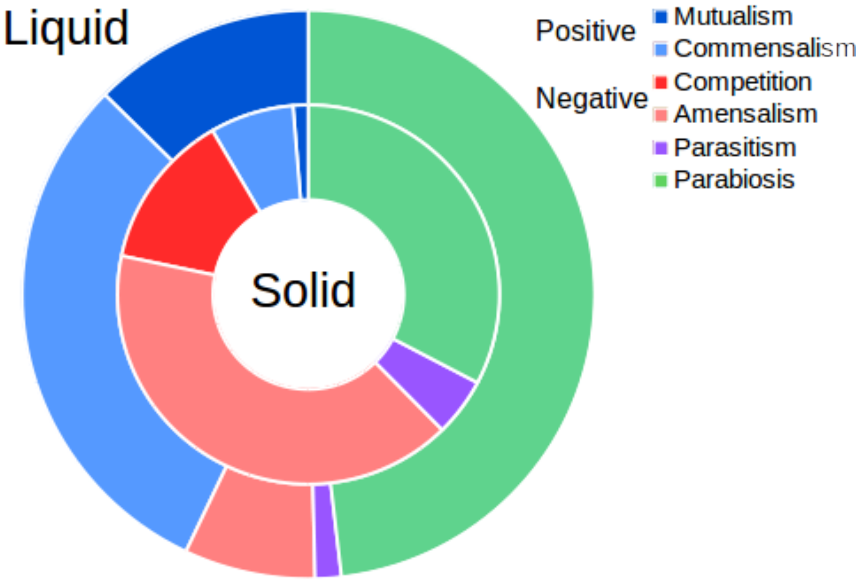


**Figure S17.** Positive and negative interactions between kefir species in milk (liquid) and on milk plates (solid).

*L. lactis* background no background

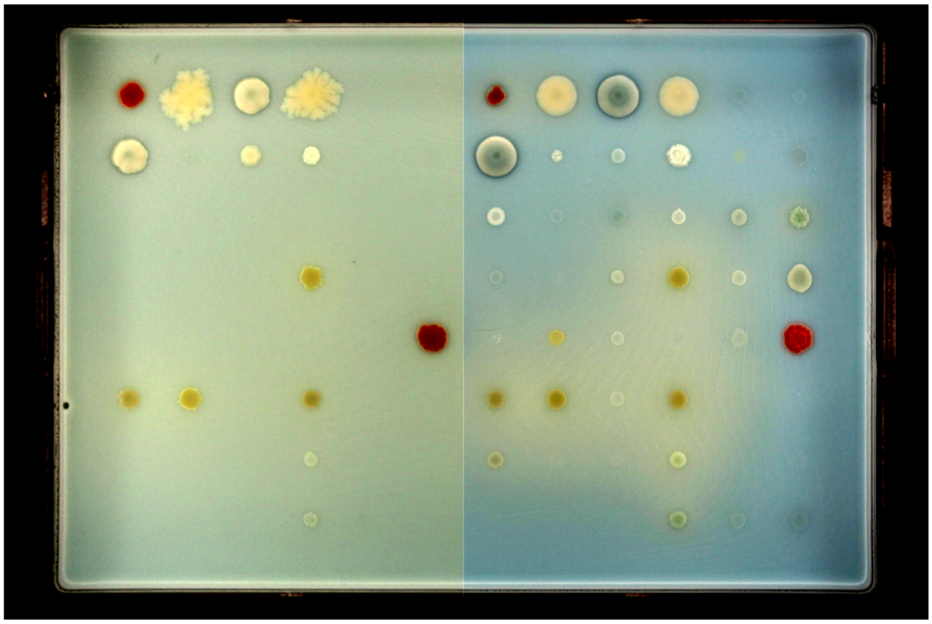

**Figure S18.** *L. lactis* is a potent inhibitor of other species when plated as a background lawn on milk plates (80 % milk, 20 % water agar and bromocresol green). Left panel: *L. lactis* background, right panel: no background species.

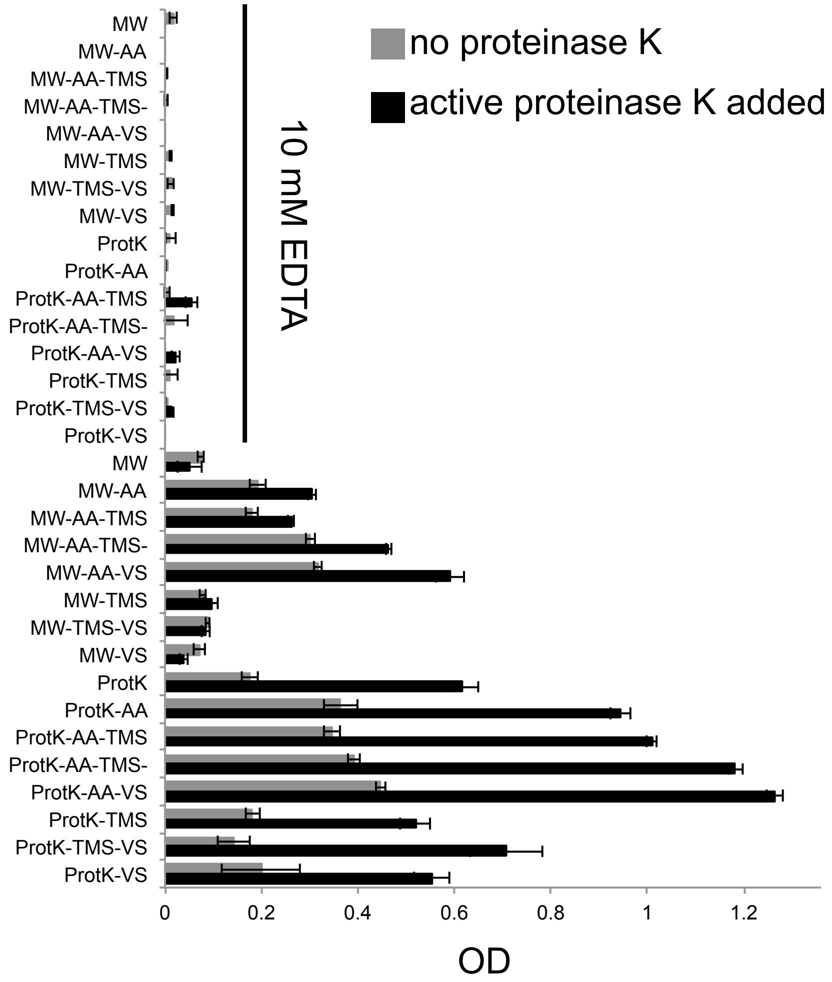


**Figure S19.** *L. mesenteroides* monoculture growth with and without proteinase K (0.2 mg/ml) and EDTA. Milk whey medium (MW) and protein rich whey medium (ProtK) was supplemented with different combinations of trace metals (TMS), vitamins (VS) and amino acids (AA).

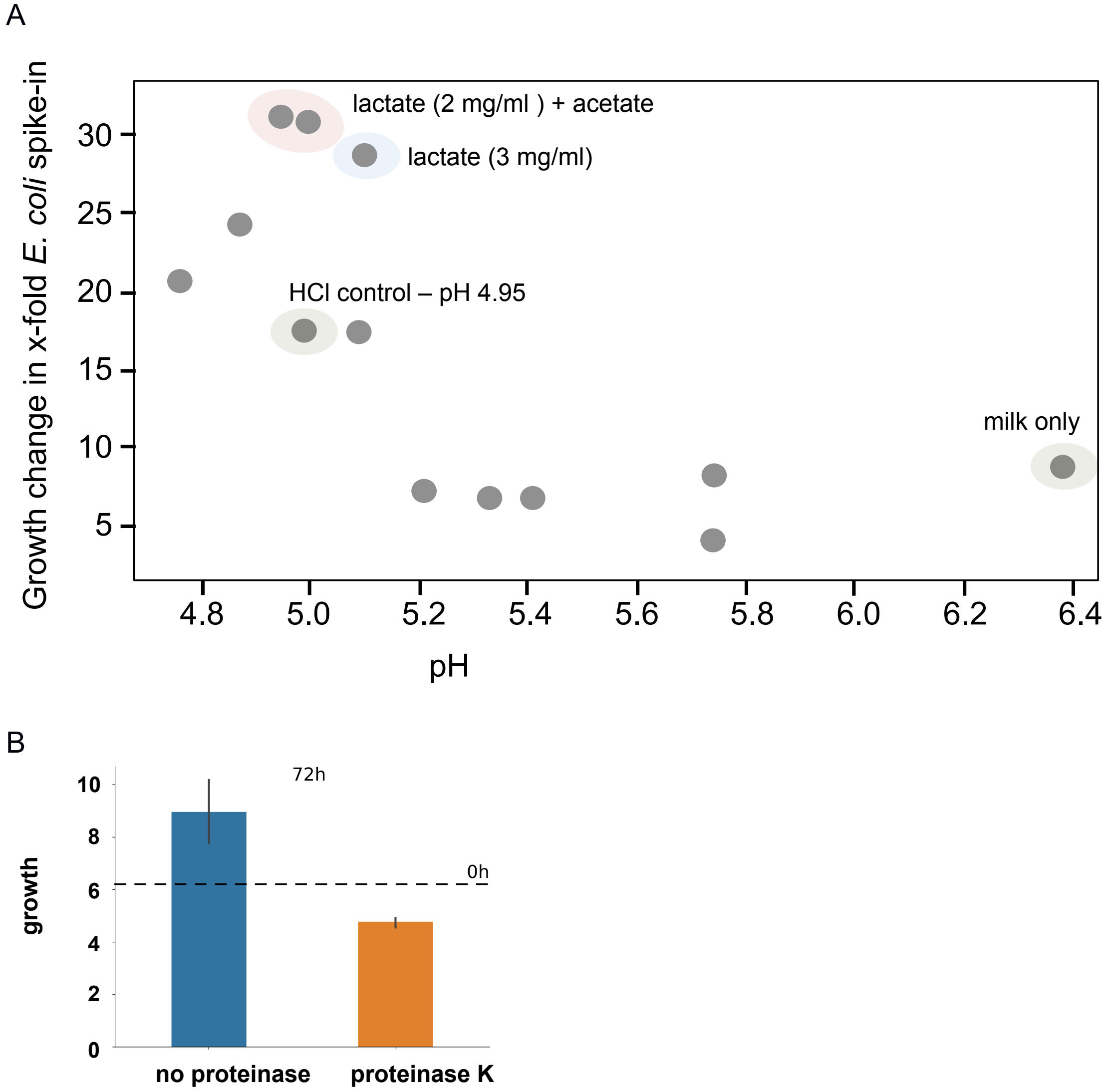


**Figure S20.** *L. kefiranofaciens* profits from lactate and acetate when added to milk. (A) *L. kefiranofaciens* growth in milk after 72 hours plotted against pH. There is a clear preference for growth around pH 5, but it cannot explain the effect fully, since pH adjustment with HCl or acetate does not result in comparable growth as lactate containing samples. (B) Addition of proteinase K to milk had no effect on *L. kefiranofaciens* growth.

Untargeted FIA-TOF

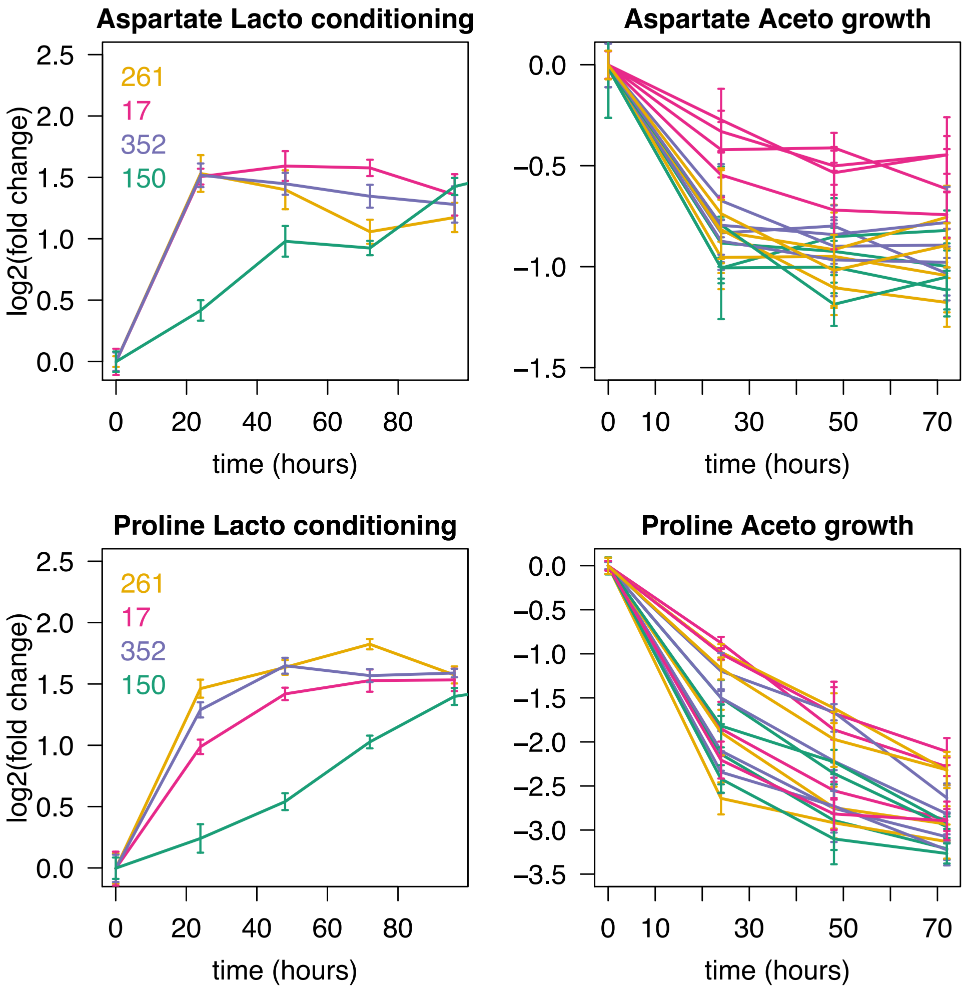


**Figure S21.** Aspartate and proline consumption by *Acetobacter* in *L. lactis* spent medium. Left penal: Milk was conditioned by four different *L. lactis* strains isolated from kefir (SB-17, SB-150, SB-261 and SB-352). Increase in aspartate and proline was measured every 24 hours by FIA-qTOF MS. Right panel: Four different *Acetobacter* strains (*A. fabarum*: SB-290, SB-373 and *A. ghanensis*: SB-354, SB-380) isolated from kefir were grown in spent whey and aspartate and proline consumption was measured by FIA-qTOF MS after 24, 48 and 72 hours.

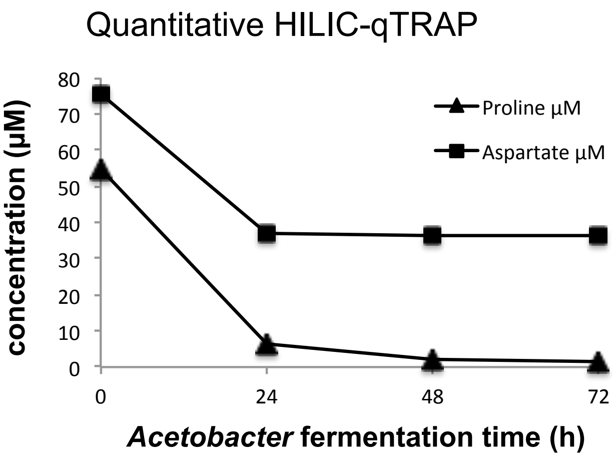


**Figure S22.** Quantification of aspartate and proline consumption by *A. fabarum* in *L. lactis* spent whey.

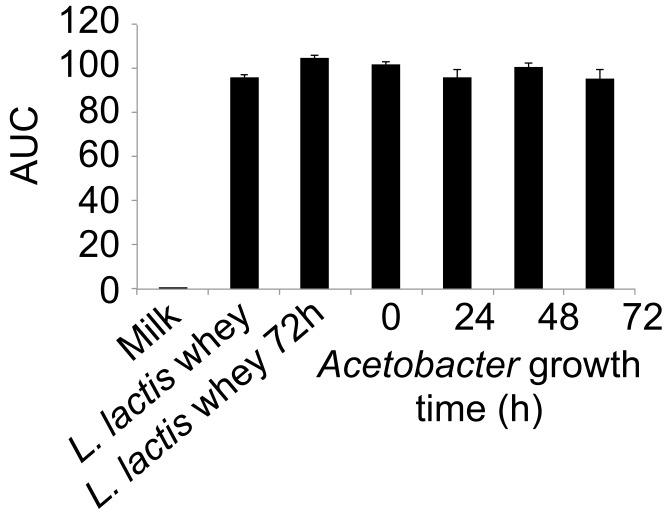


**Figure S23.** Lactate concentration in spent whey after *L. lactis* fermentation and during *A. fabarum* growth in *L. lactis* spent whey. *A. fabarum* does not consume considerable amounts of lactate during 72 hours of growth in spent whey.

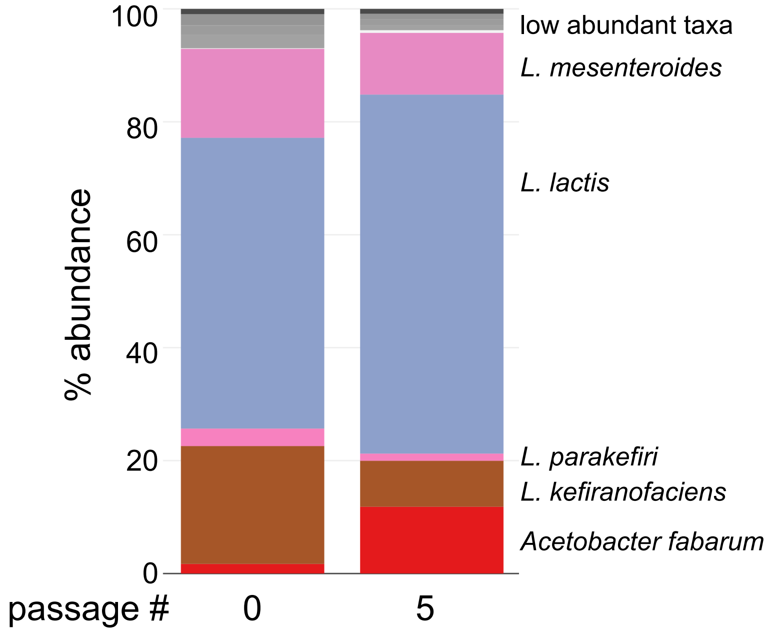


**Figure S24.** Kefir community passage using kefir fermented milk as inoculum instead of the kefir grain.
